# Reversion mutations define minimal *BRCA1/2* requirements for therapy resistance

**DOI:** 10.64898/2026.05.22.727163

**Authors:** Lorena Magraner-Pardo, William Kerrison, Dragomir B. Krastev, Rachel Alcraft, Hui Xiao, Rachel Brough, Feifei Song, Sheyne Choi, Aditi Gulati, Manuel Rodrigues, Intidhar Labidi-Galy, Eric Pujade-Lauraine, Isabelle Ray-Coquard, Syed Haider, Stephen J. Pettitt, Andrew N. J. Tutt, Christopher J. Lord

**Affiliations:** Precision Oncology Laboratory, The Breast Cancer Now Toby Robins Research Centre, The Institute of Cancer Research, London, SW3 6JB, UK; Scientific Computing, The Institute of Cancer Research, London, SW3 6JB, UK; Breast Cancer Data Science Laboratory, The Breast Cancer Now Toby Robins Research Centre, The Institute of Cancer Research, London, SW3 6JB, UK; Institut Curie, Paris, and GINECO, France; Department of Oncology, Hôpitaux Universitaires de Genève, Geneve, Switzerland; Department of Medicine, division of Oncology, Faculty of Medicine, University of Geneva, Genève, Switzerland; ARCAGY Research, Paris, France; Centre Léon Bérard, Lyon, and GINECO, France

## Abstract

Although platinum salts or PARP inhibitors are effective in delivering anti-tumor responses in people with *BRCA1* or *BRCA2* mutated cancers, drug resistance is common and is often caused by secondary *BRCA1/2* reversion mutations that restore function. By collating and analyzing 848 *BRCA1/2* reversion mutations in 384 cancer patients with drug resistance, we confirm that pathogenic *BRCA1/2* mutation type influences the acquisition of reversions, and that large *BRCA1/2* deletions are an underappreciated form of reversion. Integrating reversion data with systematic CRISPR-Cas9 screens that delete *BRCA1/2* exons, we also show that both proteins contain privileged domains whose structure is essential for drug resistance, including the PALB2 interacting domains of both BRCA1 and BRCA2. Reversions in PALB2 also conserve both BRCA1 and BRCA2 binding domains. Surprisingly, exon 11 of BRCA2, which encodes BRC repeats 1-8, is not essential for resistance. Using this patient and functional information, we estimate the likelihood of pathogenic *BRCA2* mutations to revert. We show that risk of reversion correlates with both the presence of clinical reversions and the response to treatment, suggesting that the propensity to revert could be a useful clinical parameter.

## Main Text

Platinum salt-based chemotherapy or poly(ADP-ribose) polymerase (PARP) inhibitors (PARPi) (*1, 2*) are now routinely used to treat homologous recombination defective (HRD) cancers, including those with pathogenic, loss-of-function mutations in *BRCA1* or *BRCA2 (BRCAm)* (*3*). Although PARPi and platinum salts deliver significant and sustained anti-tumor responses in *BRCAm* cancer patients, resistance to these agents often occurs (*4–6*). A recurrently occurring mechanism of PARPi resistance are secondary reversion mutations in either *BRCA1* or *BRCA2*, i.e. secondary somatic mutations that restore the native open reading frame (*7–10*). *BRCA1/2* reversions are common in each of the four cancers where PARPi are routinely used: breast, ovary, prostate and pancreatic cancers (*9, 10*). For example, 57% of advanced *BRCAm* breast cancer patients have reversion mutations at the point of platinum salt or PARPi resistance (*5*), whereas in *BRCA*m metastatic castration resistance prostate cancer, 80% of patients have reversions at the point of PARPi resistance (*4*). Importantly, the presence of *BRCA1/2* reversions in cancer patients prior to the use of PARPi is associated with significantly shorter time to progression (*5, 11*), suggesting that the detection of reversions could be important in predicting the overall efficacy of treatment. Different forms of *BRCA1/2* reversion have been described. For example, nonsense (stop-gain) pathogenic *BRCA1/2* mutations can be directly reverted to encode an amino acid through a secondary single nucleotide change, whereas frameshift pathogenic *BRCA*m can revert either by compensating insertion/deletion mutations that restore the native open reading frame, or via in-frame deletions that remove the pathogenic mutation (Figure 1A). In each of the above scenarios, the reverted *BRCA1/2* gene is predicted to encode a near to full-length reverted protein sufficient to mediate DNA repair (*9, 10*).

**Fig. 1.**
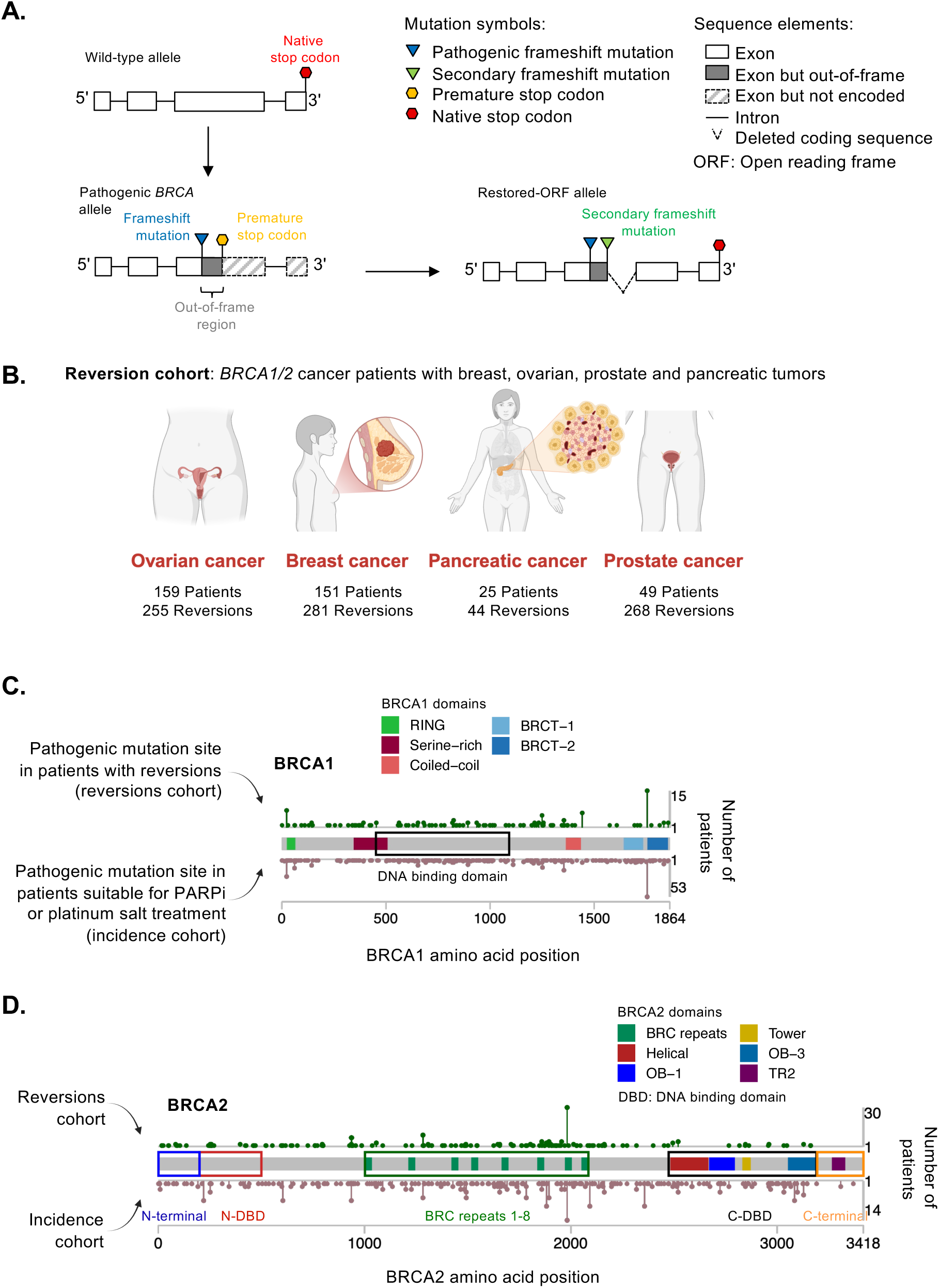
*BRCA1/2* reversion mutations associated with resistance to HRD targeting therapies in cancer. (**A**) Schematic illustrating how frameshift-causing *BRCA1/2* pathogenic mutations inactivate *BRCA1/2* and reversion mutations (in this case a secondary frameshift mutation) restore *BRCA1/2* open reading frame. **(B)** *BRCA1/2* reversion mutations in 384 *BRCA*m breast, ovarian, prostate or pancreatic cancer patients with PARPi and/or platinum salt resistance. **(C, D)** Pathogenic *BRCA1/2* mutations in reversion and incidence cohorts. Schematic shows the position within the BRCA1/2 protein structures of pathogenic mutations in the 384 patients with *BRCA1/2* reversion mutations (shown above the protein, dark green color) compared to the position of pathogenic mutations in a population of *BRCA*m patients considered for platinum or PARPi therapy - the incidence cohort (n= 1018, shown below the protein, brown color). Known functional domains within the BRCA1 and BRCA2 proteins are highlighted.

Description of clinical reversion mutations has been somewhat piecemeal, relying on initial studies in tumor cell lines (*7, 8*), followed by small studies in patient cohorts (*10*), the largest so far involving 91 patients (*9*). Although these studies were relatively small, they did suggest that: (i) most reversion mutations are small indels close to the pathogenic mutation, possibly reflecting a fitness advantage for near-full length revertant proteins (as in Fig. 1A); and (ii) that *BRCA2* pathogenic mutations in the extreme C-terminus might be less likely to revert than those elsewhere in the gene (*9*). These preliminary findings led us to now considerably expand the analysis of clinical reversions. We did this on the basis that an expanded analysis could allow us to define the domains of the BRCA1 or BRCA2 proteins that are critical to mediate treatment resistance in patients. Such analysis could also allow us to assess whether there are pathogenic *BRCAm* that are more or less likely to revert, information that could ultimately be used to inform how cancer patients are best treated.

In recent years, three factors have facilitated a greater ability to detect and then use reversion mutations to understand *BRCA1/2* function: (i) the introduction of multiple different PARPi, as well as platinum salts, into the routine treatment of *BRCA*m cancers; (ii) the consequent increase in the number of patients with clinical resistance to either drug class; and (iii) the wider use of sequencing capture panels that allow sequencing of *BRCA1/2* intronic as well as exonic sequences from tumors and circulating tumor DNA (ctDNA) (*5*), a technological improvement that enhances the ability to detect reversion mutations. Exploiting these factors, we collected and analyzed 848 reversion mutations from 384 *BRCA*m cancer patients with resistance to either PARPi and/or platinum salts. By systematically analyzing reversion mutations, including large *BRCA1/2* deletions that appear to be a poorly recognized reversion form, we find that BRCA1 and BRCA2 contain privileged domains whose structure is retained in reversions and that PARPi resistance can be mediated by reverted *BRCA2* forms that lack exon 11 encoded BRC repeats. Using the information gained from a systematic analysis of reversions, we also developed a predictor of the risk that a pathogenic *BRCA2* mutation reverts, showing this property of *BRCA2* mutations is associated with clinical outcome.

## Results

### Collation of a cancer patient reversions dataset

We collated a patient cohort of 384 *BRCA*m cancer patients with either platinum salt and/or PARPi resistance who had acquired *BRCA1* or *BRCA2* reversion mutations (termed forthwith, the “reversions cohort”, Fig. 1B) derived from 59 clinical studies (Fig. S1, Data S1 and S2). In this reversions cohort, 79% of patients had a unique pathogenic *BRCA*m. Most patients (71.4%) had a single reversion mutation, with the remainder exhibiting more than one reversion event (2-34 reversions, Fig. S2A), giving us a final total of 848 *BRCA1/2* reversion mutations (Data S3). Most patients with multiple reversions had been profiled using ctDNA (described in detail in Fig. S2B-G), which has an increased ability to detect reversions over solid tumour sequencing. Most *BRCA1/2* reversions were deletions flanked by microhomology sequences (1-8 nucleotides, Fig. S3, Data S4, 5), an observation consistent with the previous proposition that error-prone DNA repair mechanisms that require regions of microhomologous sequence, including Theta Mediated End Joining (TMEJ), are a major contributor to reversion formation (*9, 12*).

### Pathogenic *BRCA1/2* mutation type influences the acquisition of reversions

To explore the propensity of different pathogenic mutation types to revert, we compared the pathogenic *BRCA1/2* mutations seen in the reversions cohort (Data S6) to those seen more generally in *BRCA*m patients who would be considered for platinum or PARPi therapy (i.e. an “incidence cohort”, Fig. 1C, D, S7, S8; *Supplementary Methods*). This comparison confirmed two linked observations (*9*), namely that, when compared to their frequency in the incidence cohort, reversion of pathogenic frameshift or stop-gain mutations appeared to be overrepresented in the reversions cohort, whereas reversion of pathogenic *BRCA1/2* splice site or missense mutations were significantly underrepresented events (Fig. 2A, Data S9), suggesting that pathogenic frameshift or stop-gain pathogenic mutations have a greater propensity to revert. This is consistent with the idea that there might be a more restricted set of ways in which function can be restored from some pathogenic mutation types (such as splice site or missense mutations), when compared to others (such as frameshifts). This hypothesis appeared to be supported when we examined the varied ways that different types of pathogenic mutation reverted (Fig. 2B). Stop gain mutations tended to revert via two routes: secondary substitutions that restored an amino acid codon or via in-frame, intra-exon deletions (Fig. 2B). In contrast, frameshift pathogenic mutations reverted via multiple routes (Fig. 2B), each of which restored the open reading frame (ORF): most frequently the restoration of the ORF was achieved by an intra-exon deletion that spanned the pathogenic mutation, or via a compensating secondary frameshift mutation (SFM) occurring between the pathogenic mutation and an upstream or downstream premature out-of-frame stop codon. SFMs usually result in the retention of out-of-frame protein sequence between the pathogenic and reversion mutation, effectively losing the wild type protein sequence in this region. Thus, reversion of frameshift mutations is overall more likely to result in loss of sequence than stop-gains. Bypass of either type of pathogenic mutation by mutations resulting in alternative splicing has also been described, which could effectively delete entire exons from the mutated protein (*6, 9*).

**Fig. 2.**
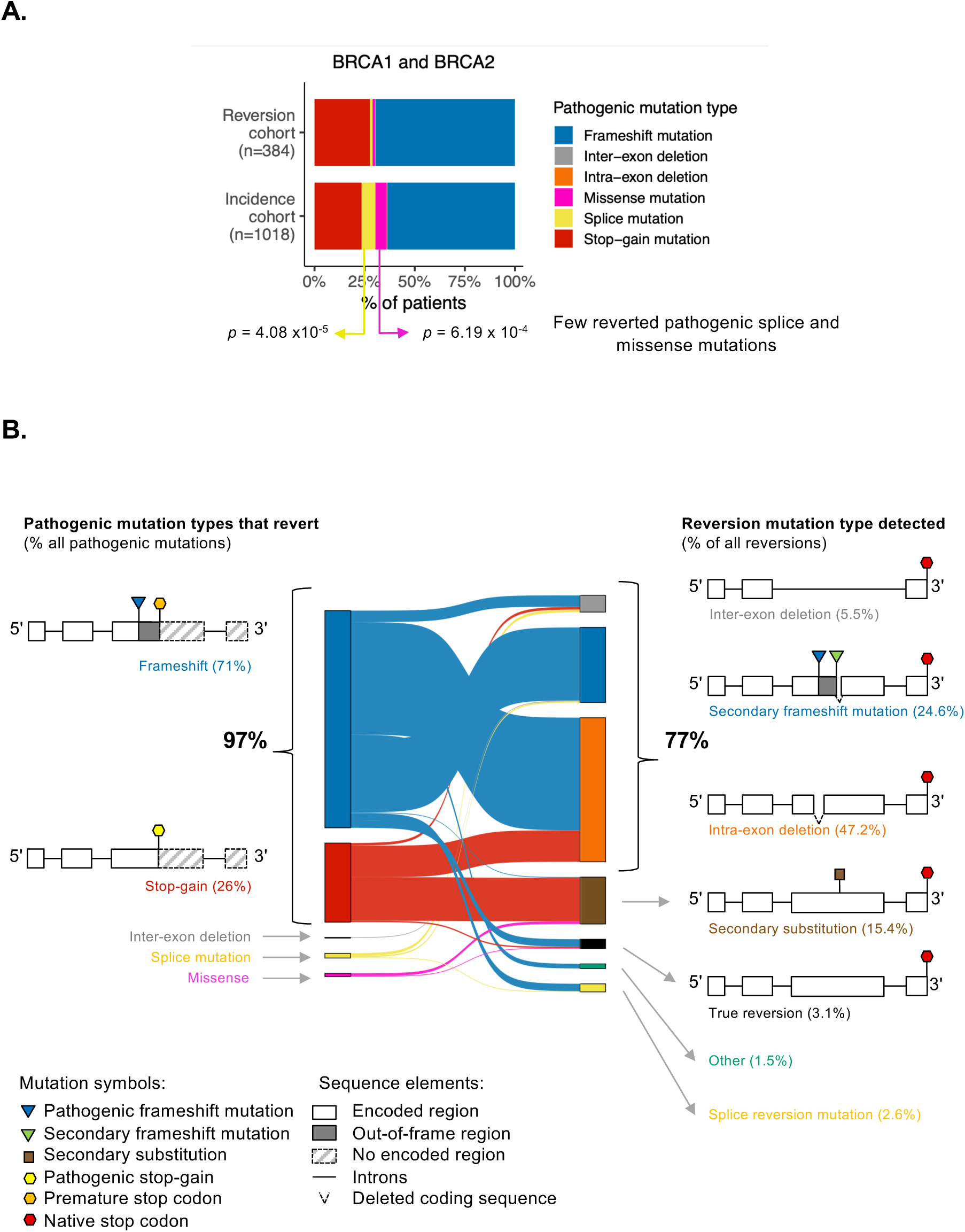
Comparison of *BRCA1/2* mutations in the reversions *vs*. incidence cohorts. **(A)** Column chart illustrating a comparison between reversions and incidence data for different pathogenic mutation types is shown. Reversions in splice site (*p* = 4.08 x10^−5^) or missense pathogenic mutations (*p* = 6.19 x10^−4^) are underrepresented in the reversions cohort compared to the incidence cohort. *p* values calculated by Fisher’s exact adjusted for multiple testing using the Benjamini-Hochberg method. (**B**) Most *BRCA1/2* reversion mutations result in loss or disruption of protein sequence. Sankey plot depicting the distribution of pathogenic mutation types (left) that give rise to reversion mutations, along with the corresponding reversion mutation types (right), scaled by their occurrence in the reversion cohort. The majority of reversions are derived from pathogenic frameshift and stop-gain mutations (97%). 77 % of reversions result in loss or disruption of > 1 amino acid, reverting via either secondary frameshift mutations, intra-exon deletions, or inter-exon deletions which span the pathogenic mutation site.

### *BRCA1/2* deletion-mediated reversions define the regions of BRCA1/2 sufficient for treatment resistance

In a prior analysis of clinical reversion mutations, large deletions (i.e. those over 2 kb) in *BRCA1* or *BRCA2* had not been reported (*9*). However, when we analyzed the larger patient cohort described here, deletions involving introns as well as partial or complete exons, or even multiple exons, were observed in 5.5% of patients (inter-exon deletion, Fig. 2B). For example, in case KCL644, a deletion reversion (KCL644_2) was present in *BRCA2* that deleted DNA from a site in intron 10 through to a site in the coding sequence in exon 11, deleting at least 1274 amino acids from the predicted protein as well as likely affecting splicing (Fig. 3A, B). Similarly, large deletion reversions were seen in *BRCA1*, such as in case KCL725, where two reversions were identified (KCL725_2 and KCL725_4) that removed coding sequence for over 600 amino acids (Fig. 3C, D). As well as improvements in mutation calling, the ability to identify larger *BRCA1/2* deletion reversions such as these might be caused by the increased use of sequencing capture panels that cover both intronic as well as exonic *BRCA1/2* sequence, thus allowing deletion break points within introns to be identified (*5, 13*).

**Fig. 3.**
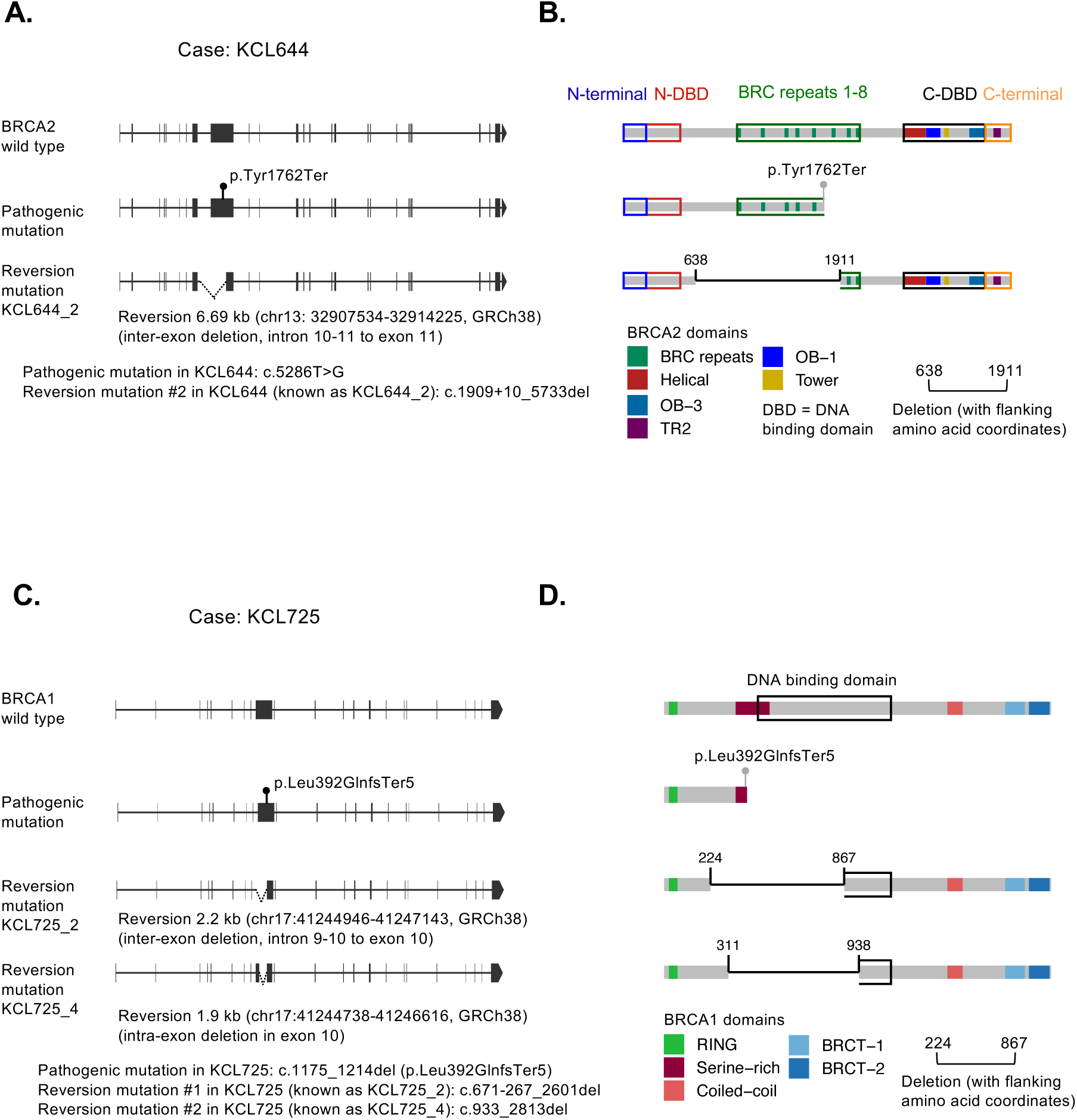
Inter-exon deletions in *BRCA1/2.* (**A**) Example of an inter-exon deletion involving multiple *BRCA1* exons in case KCL644. *BRCA2* intron/exon structure for wild type *BRCA2* is shown above that for the pathogenic mutation in case KCL644 (*BRCA2*:c.5286T>G, p.Tyr1762Ter) and a large deletion reversion (KCL644_2 *BRCA2*: c.1910_5733del) that encompasses all of the intron between exon 10 and 11 and part of exon 11. By partially deleting exon 11 and its splice acceptor, the reversion in KCL644 is predicted to facilitate an in-frame splicing event between exons 10 and 12. (**B**) Predicted protein domain structures for wild type BRCA2, BRCA2 encoded by the KCL644 pathogenic mutation (which truncates the protein at amino acid 1762) and the KCL644_2 reversion mutation. The KCL644_2 reversion is predicted to facilitate deletion of 1273 amino acids between residues 638 and 1911. (**C**) Example of an inter-exon deletions involving *BRCA1* exonic region as well as intronic sequence in case KCL725. BRCA1 intron/exon structure for wild type *BRCA1* is shown, above that for the pathogenic mutation in case KCL725 (*BRCA1*: c.1175_1214del, p.Leu392GlnfsTer5) and two different deletion reversions in KCL725 (KCL725_2 *BRCA1*: c.671-267_2601del and KCL725_4 c.933_2813del). (**D**) Predicted protein domain structures for wild type BRCA1, BRCA1 encoded by the KCL725 pathogenic mutation and the reversion mutations KCL725_2 and KCL725_4.

As the majority of reversion mutations (77 %) removed part of the coding sequence (i.e. intra-exon deletions, plus SFMs and inter-exon deletions, Fig. 2B), we asked whether these events were limited to certain regions or functional domains in the BRCA1 or BRCA2 proteins that may be therefore classed as being less important for therapeutic resistance, and thus more permissive to be completely deleted or disrupted by reversion mutations (e.g. via deletion, substitution, or replaced by out-of-frame sequences generated by secondary frameshift reversions). We reasoned that by mapping all of the regions of BRCA1/2 that were disrupted by reversion mutations, we might be able to infer the propensity of different pathogenic mutations to revert and also infer which domains of BRCA1 and BRCA2 are sufficient for clinical resistance to treatment. To assess this, we took two approaches: (i) mapping the domains of BRCA1 or BRCA2 that were predicted to be either completely or partially removed from wild-type proteins by reversions that were deletions (Figure 4 A,C); and (ii) using the frequency with which each amino acid in BRCA1 or BRCA2 was either deleted or changed by a reversion mutation to estimate whether it was critical or not for therapeutic resistance (described in Fig. 5A-C). As described later (Figs. 6 and 7), we then complemented this analysis by functionally assessing whether *BRCA2* alleles lacking potentially critical regions could cause cellular PARPi resistance.

**Fig. 4.**
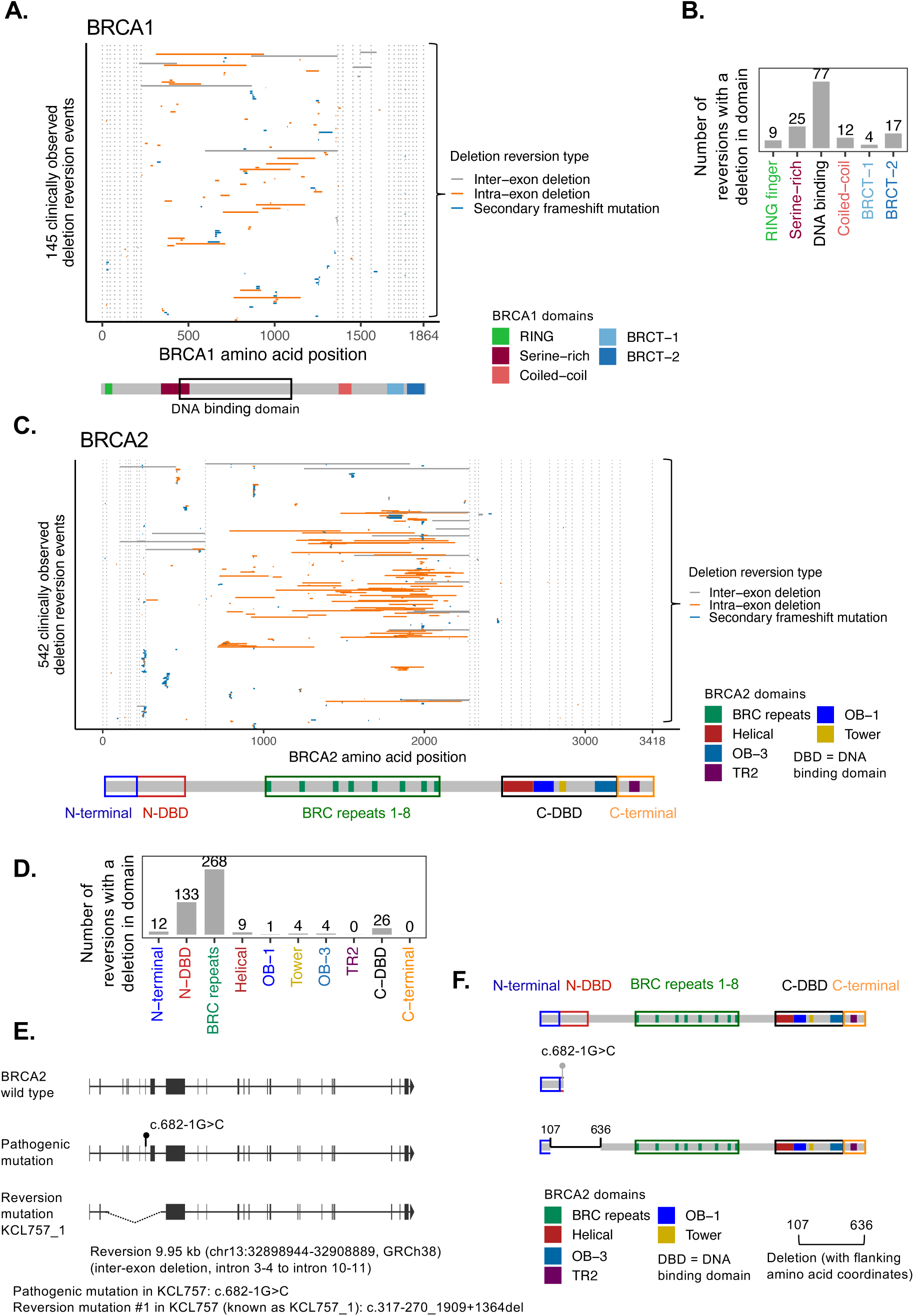
Deletion reversions in *BRCA1* and *BRCA2* that delete coding sequence. **(A)** Alignment of *BRCA1* deletion reversions in the reversions cohort with BRCA1 amino acid positions. Deletions (horizontal lines) depict the three types of deletion reversions: Inter-exon deletion (grey), intra-exon deletion (orange), and secondary frameshift mutations (blue). Vertical dotted lines indicate exon boundaries. Below the plot, BRCA1 protein domains are displayed. Most deletion reversions cluster in the central region of the BRCA1 protein. (**B**) Column chart summarizing the frequency with which each BRCA1 domains is disrupted by deletion. Most deletion reversions are in the DNA binding domain. **(C)** Alignment of *BRCA2* deletion reversions in the reversions cohort with BRCA2 amino acid positions. Key as for (A). (**D**) Column chart summarizing the frequency with which each BRCA2 domain is disrupted by deletion. Most deletion reversions cluster around the central portion of BRCA2 encompassing BRC repeats 1-8. (**E**) Example of an inter-exon deletion involving multiple *BRCA2* exons in case KCL757. *BRCA2* genomic intron/exon structure for wild type *BRCA2* is shown, above that for the pathogenic mutation in case KCL757 (*BRCA2*: c.682-1G>C in a splice acceptor site 5’ of exon 9) and a deletion reversion (KCL757_1, *BRCA2*: c.317-270_1909+1364del) that encompasses exons 4-10 which is predicted to delete the pathogenic mutation and result in an in-frame splicing event between exons 3 to 11. (**F**) Predicted protein domain structures for wild type BRCA2, BRCA2 encoded by the KCL757 pathogenic mutation (c.682-1G>C) and the reversion mutation in KCL757.

**Fig. 5.**
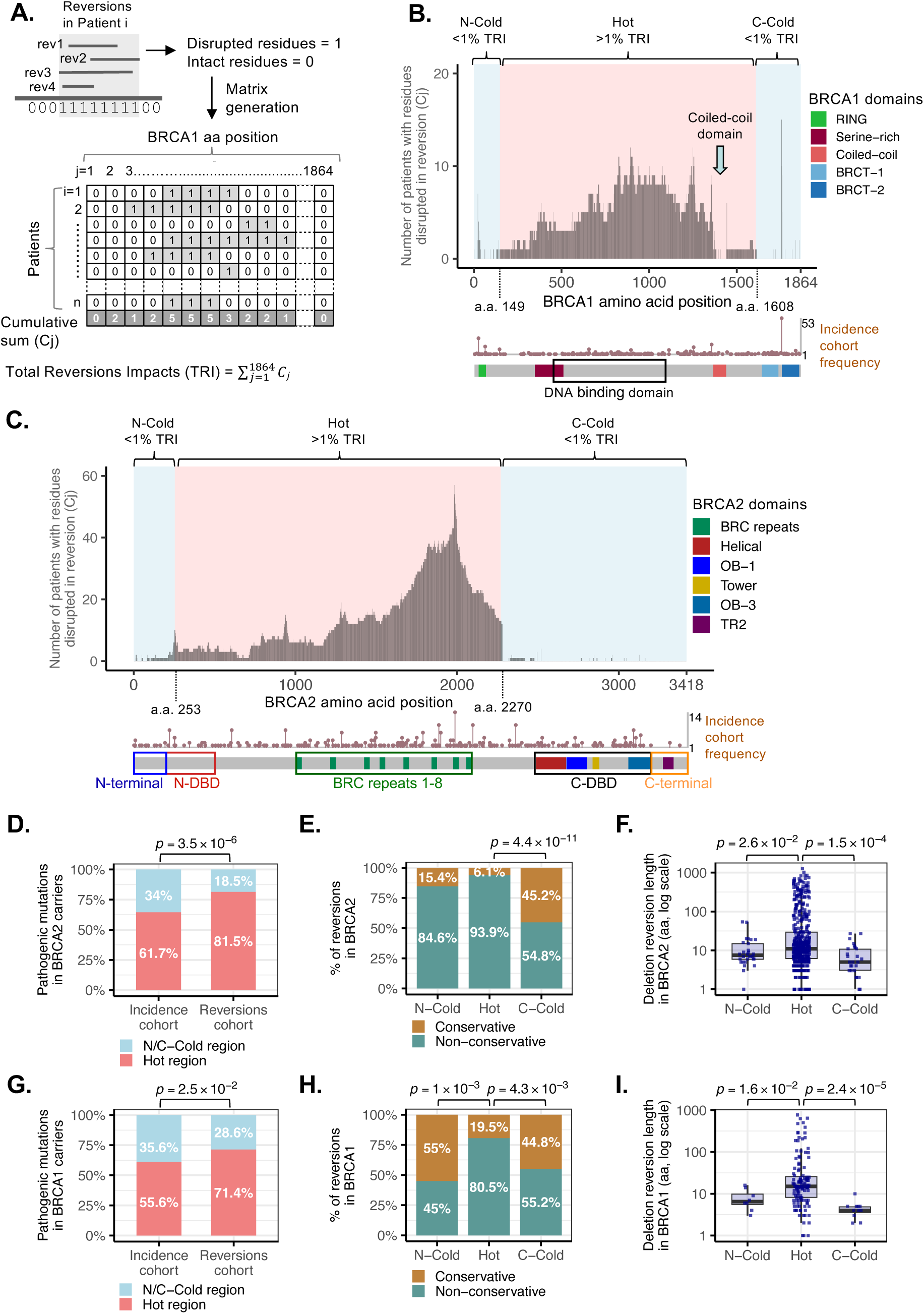
Amino acid residue-by-residue analysis of reversion mutations in BRCA1 and BRCA2. **(A)** Schematic of matrix generated to assess the frequency at which each residue in BRCA1 is disrupted by reversion (residue-by-residue analysis). Rows correspond to patients, and columns to amino acid positions in BRCA1. Amino acid residues disrupted by reversions (deletion, substitution, or frameshifted sequence) were marked as 1, whilst those not altered by reversions were annotated as 0. The matrix has one row per patient; when multiple reversions occurred in a single patient, these were collapsed, with each disrupted residue being counted only once. The cumulative sum of alterations per residue (*C_j_*, bottom row) was used to map the reversion density profile across the protein. The total reversion impact (TRI) was defined as the sum of *C_j_* across the whole protein. The same strategy was followed to generate a matrix for BRCA2 reverted cancers. **(B, C)** Per-residue map of BRCA1 (B) or BRCA2 (C) amino acids altered by reversion mutations. The histogram (grey) indicates the number of patients in which each amino acid is disrupted by reversions across the reversion cohort (*C_j_*). Regions in the N- and C-terminal regions of either BRCA1 or BRCA2 rarely altered by reversions (i.e. cumulative TRI < 1%) were defined as N- and C-Cold regions for reversions (blue area). Conversely, regions in BRCA1 or BRCA2 commonly altered by reversions (cumulative TRI >1%) were defined as “hot” regions (central pink area). For BRCA1, cold regions were defined as amino acid residues 1-149 (N-Cold) and 1608-1864 (C-Cold), and the hot region as the central segment from amino acid residues 150 to 1607. For BRCA2, cold regions were defined as residues 1-253 (N-Cold) and 2270-3418 (C-Cold), and the hot region defined as the central segment from amino acid residue 254 to 2269. **(D)** Column chart showing that pathogenic BRCA2 mutations in cold regions are significantly underrepresented in the reversions cohort compared to the incidence dataset (Fisher’s exact test, *p* = 3.5 x10^-6^). **(E)** Column chart showing that BRCA2 reversions in cold regions have a higher proportion of conservative reversions (e.g. substitutions or true reversions), when compared to those found the hot region (Fisher’s exact tests *p* = 8.1 x10^-2^ and 4.4 x10^-11^). **(F)** Mean deletion reversion length is reduced in N-Cold (Wilcoxon rank-sum test *p* = 2.6 x 10^-2^, left) and C-Cold (Wilcoxon rank-sum test *p* = 1.5 x 10^-4^, right) regions of BRCA2, compared to the hot central region (central boxplot), suggesting that reversion events in cold regions are more constrained. **(G)** Column chart showing that pathogenic BRCA1 mutations in cold regions are significantly underrepresented in the reversions cohort compared to the incidence dataset (Fisher’s exact test, *p* = 2.5 x10^-2^). **(H)** Column chart showing that BRCA1 reversions in cold regions have a higher proportion of conservative reversions (e.g. substitutions or true reversions), when compared to those found the hot region (Fisher’s exact tests *p* = 1.1 x10^-3^ and 4.3 x10^-3^). **(I)** Mean deletion reversion length is reduced in N-Cold (Wilcoxon rank-sum test *p* = 1.6 x 10^-2^, left) and C-Cold (Wilcoxon rank-sum test *p* = 1.4 x 10^-5^, right) regions of BRCA1, compared to the hot central region (central boxplot), suggesting that reversion events in cold regions are more constrained.

**Fig. 6.**
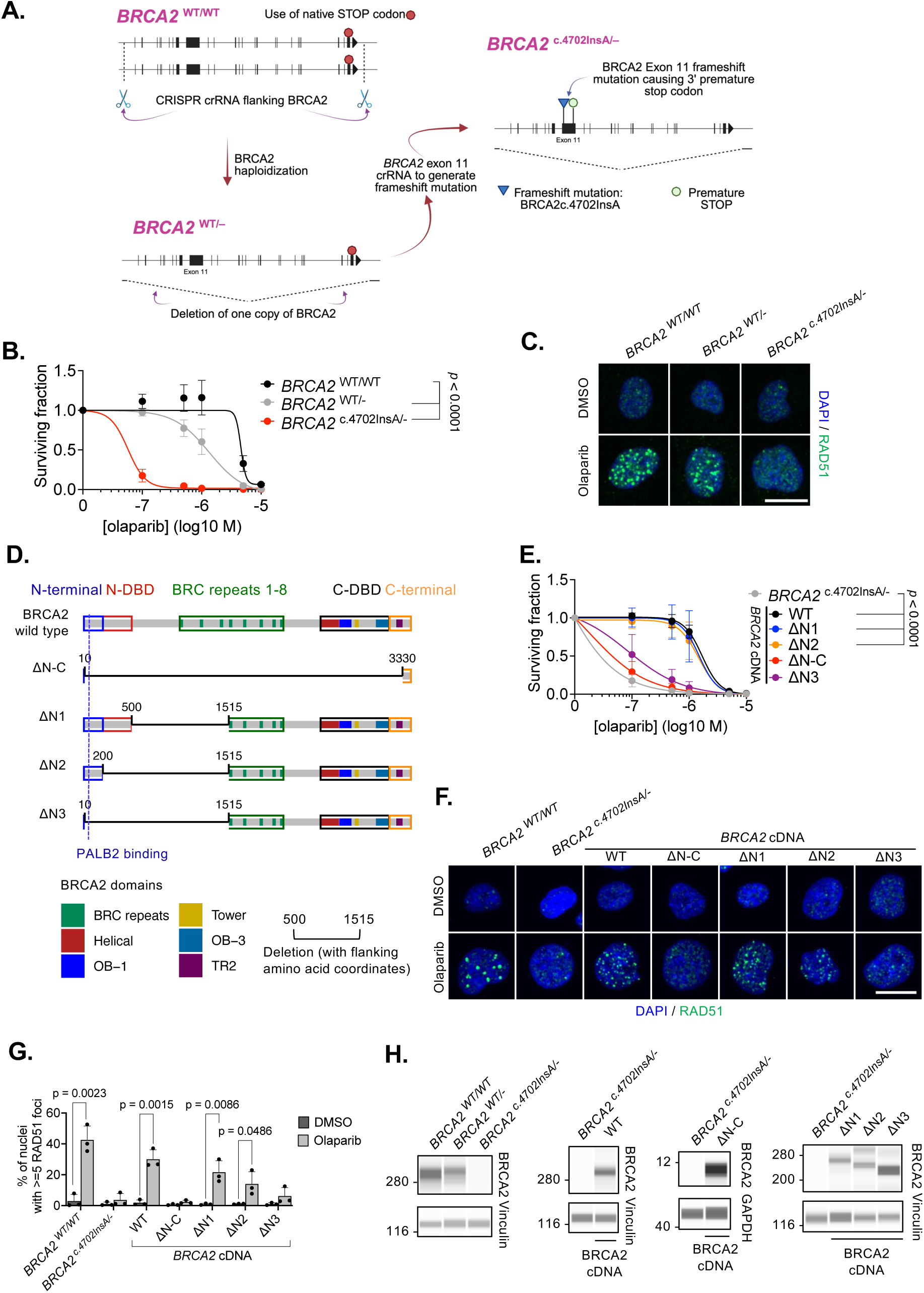
*BRCA2* transgene variants identifies domains essential for PARPi resistance. **(A)** Schematic describing the derivation of *BRCA2* mutant cell lines. CRISPR-Cas9 was used to haploidize the *BRCA2* allele in p53 mutant RPE1 cells, generating *BRCA2* ^WT/–^ cells. Subsequent mutagenesis of the remining (wild-type) *BRCA2* allele caused the formation of a clone with a premature truncation within *BRCA2* exon 11, *BRCA2* ^c.4702InsA/–^. **(B)** PARPi dose-response curves in *BRCA2*^WT/WT^, *BRCA2* ^WT/–^ and *BRCA2* ^c.4702InsA/–^ cells exposed to the PARPi olaparib for 10 days, after which survival was assessed using CellTiter Glo. Surviving fractions compared to cells exposed to the drug vehicle (DMSO) are shown. *p* values calculated using ANOVA. Error bars represent standard deviation from n = 3 experiments. **(C)** Representative images of nuclear RAD51 foci in *BRCA2*^WT/WT^, *BRCA2* ^WT/–^ and *BRCA2* ^c.4702InsA/–^ cells exposed to the PARPi olaparib (5 μM) or the drug vehicle, DMSO, for 24 hours (see Fig. S5G for quantification of RAD51 foci). Scale bar = 10 μm. Blue staining represents nuclei (DAPI), green staining represents RAD51. (**D**) Graphical representation of BRCA2 protein variants expressed via cDNA constructs in *BRCA2* ^c.4702InsA/–^ cells. ΔN1, ΔN2 and ΔN3, encode progressive deletions of the BRCA2 N-DBD and N-terminal PALB2 binding site, whereas ΔN-C has a deletion covering both N-DBD and N-terminal PALB2, BRC repeats and C-DBD. Deleted regions are indicated with a thin line and flanking amino acid co-ordinates. (**E**) PARPi dose-response curves in *BRCA2* ^c.4702InsA/–^ cells expressing cDNAs from (D) exposed to the PARPi olaparib for 10 days, after which survival was assessed using CellTiter Glo. Surviving fractions compared to cells exposed to the drug vehicle (DMSO) are shown. *p* values calculated using ANOVA. Error bars represent standard deviation from n = 3 experiments. **(F)** Representative images of nuclear RAD51 foci in *BRCA2* ^c.4702InsA/–^ cells expressing cDNAs from (D) exposed to the PARPi olaparib (5 μM) or the drug vehicle, DMSO, for 24 hours. Scale bar = 10 μm. Blue staining represents nuclei (DAPI), green staining represents RAD51. **(G)** Percentage of cells with >=5 nuclear RAD51 foci in cells exposed to either 5 µM olaparib (or drug vehicle, DMSO) for 24 hours. Error bars represent standard deviation from n = 3 experiments. *p* values calculated by Student’s t test. **(H)** Simple Western immunoblot analysis using an antibody detecting a BRCA2 C-terminal epitope in lysates generated from *BRCA2* ^c.4702InsA/–^ cells expressing BRCA2 cDNA constructs. Positions of molecular weight markers are shown on left (kDa).

**Figure 7.**
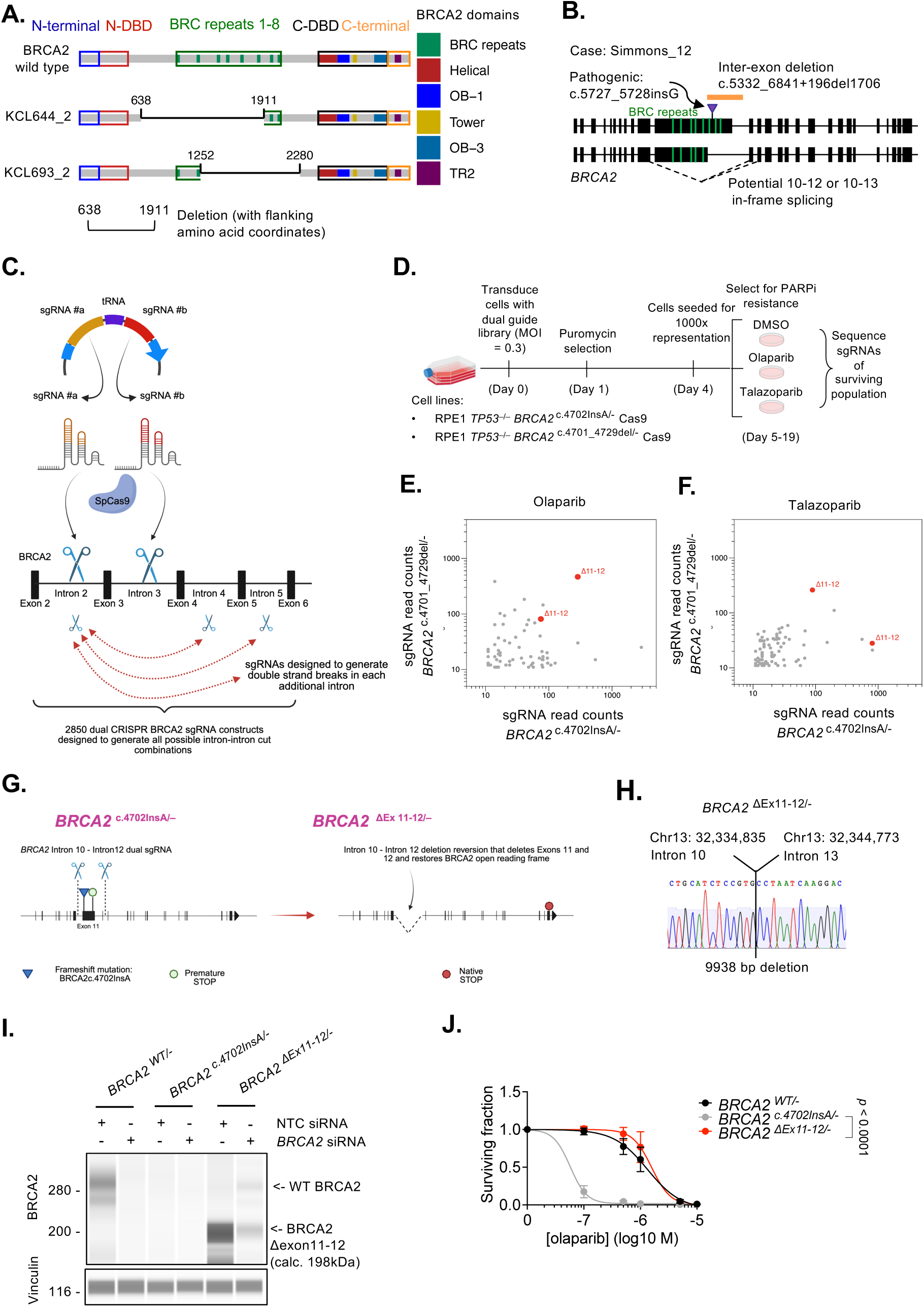
PARPi resistance can be mediated by BRCA2 forms that lack exon 11 encoded BRC repeats. **(A)** Schematic representation of the BRCA2 wild-type protein domain structure and predicted BRCA2 domain structure of reversions in cases KCL644_2 and KCL693_2, that are deletions in *BRCA2* that directly remove large parts of exon 11. **(B)** *BRCA2* intron/exon structure from a reversion case (Simmons_12) with a pathogenic *BRCA2 c.5727_5728insG* mutation within exon 11 and an inter-exon deletion reversion c.5332_6841+196del1706; the latter reversion mutation deletes the splice donor site 3’ to exon 11. This loss of the exon 11 splice donor potentially causes exon 10-12 or exon 10-13 in frame splicing events. If canonical splice sites are used, both exon 12 and exon 13 are in-frame with exon 10. **(C)** Graphical representation of dual CRISPR guide library, with pairs of guides cutting in different introns. Polycistronic sgRNA-tRNA-sgRNA expression constructs were generated to encode pairs of short guide (sg)RNAs designed to generate DNA breaks in different introns when combined with SpCas9. An example of one sgRNA pair cutting intron 2 and intron 3 is shown. A library encompassing 2850 pairs of sgRNAs targeting *BRCA2* was constructed, designed to generate all possible intron-intron break combinations as shown. **(D)** *BRCA2* dual CRISPR screen experimental workflow. Two different clones with exon 11 BRCA2 truncating mutations (*BRCA2* ^c.4702insA/–^ cells described in Figure 6, plus a second BRCA2 defective clone, *BRCA2* ^c.4701_4729del/–^) were transduced with the dual guide library and then selected using PARPi. sgRNA pairs causing PARPi resistance were identified in the surviving cell population by deep sequencing. **(E, F)** sgRNA read counts from *BRCA2* ^c.4702insA/–^cells (x-axis) and *BRCA2* ^c.4701_4729del/–^ clone (y-axis) cells following infection with *BRCA2* dual CRISPR sgRNA library followed by olaparib (E) or talazoparib (F) selection. Each point represents a sgRNA vector. Labels correspond to the exons deleted between the targeted introns, i.e. Δ11-12 corresponds to a guide pair designed to cut in intron 10 and intron 12, thus deleting both exons 11 & 12. **(G)** Schematic describing the derivation of *BRCA2* mutant cell lines with reversions that exclude exons 11 and 12. *BRCA2* ^c.4702insA/–^ cells were transduced with a dual CRISPR sgRNA construct designed to generate double strand breaks in introns 10 and 12, generating deletions that remove exon 11 truncating mutations and which facilitate in frame exon 10-exon 13 splicing, termed *BRCA2* ^ΔEx11-12/–^ cells. (**H**) DNA sequencing traces from resultant *BRCA2* ^ΔEx11-12/–^ cells. (**I**) Simple Western immunoblot analysis using an antibody detecting a BRCA2 C-terminal epitope in lysates generated from *BRCA2* ^ΔEx11-12/–^ cells. Note that full length BRCA2 migrates faster than its predicted molecular weight (384 kDa). Positions of molecular weight markers shown on left (kDa). **(J)** Olaparib dose-response curves in *BRCA2* ^ΔEx11-12/–^ cells exposed to the PARPi olaparib for 10 days, after which survival was assessed using CellTiter Glo. Surviving fractions compared to cells exposed to the drug vehicle (DMSO) are shown. *p* values calculated using ANOVA. Error bars represent standard deviation from n = 3 experiments.

**Figure 8.**
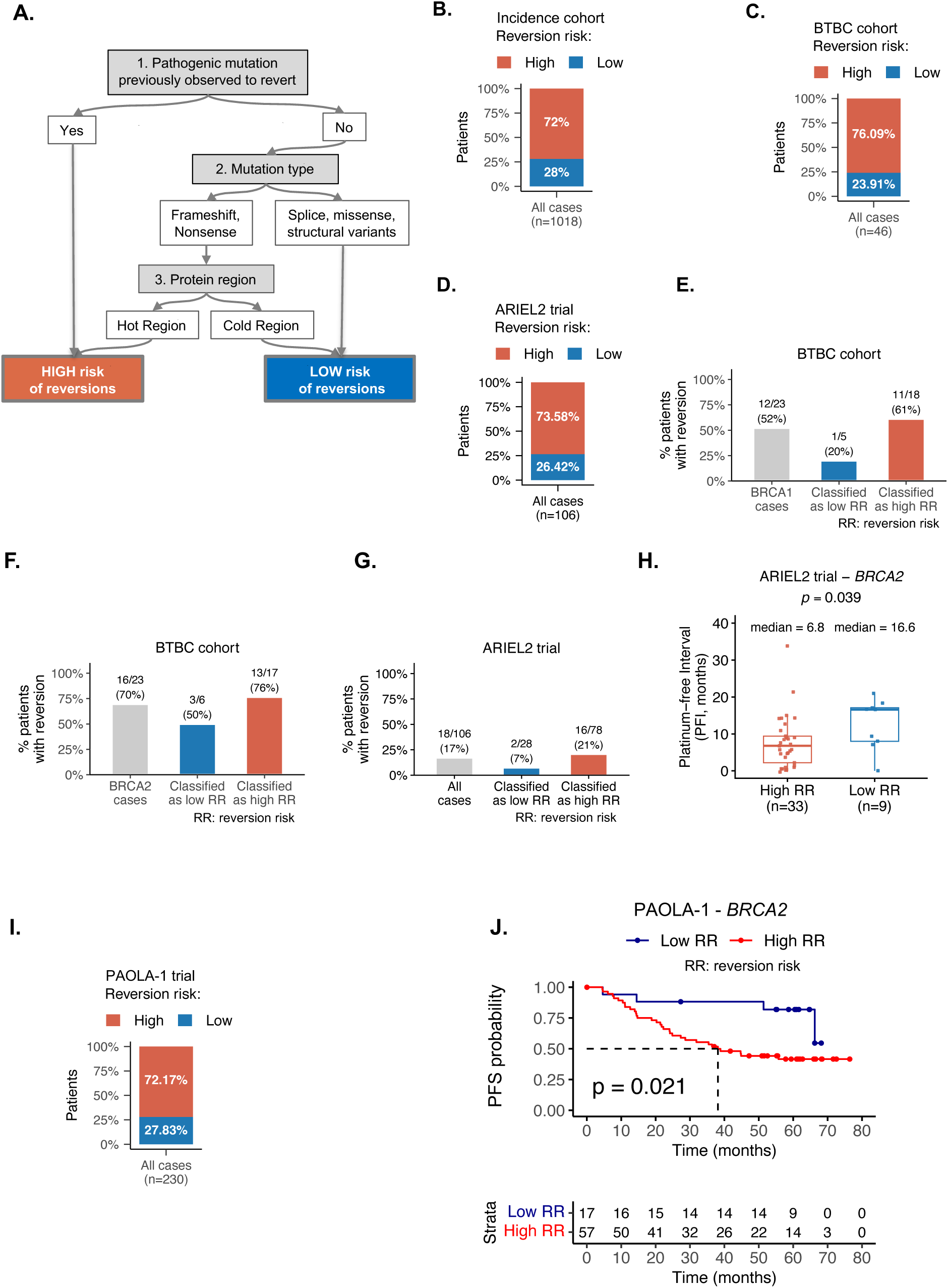
Reversion risk classifier stratifies clinical outcomes in *BRCA2* mutant cancers. **(A)** Schematic illustrating the reversion-risk algorithm designed to classify pathogenic mutations as high or low risk of reversion. The classifier uses prior evidence of pathogenic mutations being seen in the reversion cohort, the mutation type of pathogenic mutations and whether pathogenic mutations sit within “hot” or “cold” regions of *BRCA1* or *BRCA2*. **(B-D)** Frequency of patients with high (red) or low (blue) reversion risk *BRCA1/2* pathogenic mutations in the incidence cohort (B), the BTBC study (*5*) (C) or the ARIEL2 trial (*11*) (D). **(E-G)** Frequency of patients with reversions in BTBC and ARIEL2 classified according to high or low risk of reversion. Data is shown for *BRCA1* mutant patients in BTBC (E), BRCA2 mutant patients in BTBC (F) or both *BRCA1* or *BRCA2* mutant patients in ARIEL2 (G). F1 scores for each dataset are described in Data 15. **(H)** Boxplot showing the association between predicted reversion risk and platinum-free interval (PFI) in BRCA2 mutant patients in the ARIEL2 trial. Patients were stratified into high (red) or low (blue) reversion risk groups. *p* value calculated with Wilcoxon test. **(I)** Pathogenic mutations in PAOLA-1 trial (*36*) were classified as high (red) or low (blue) predicted reversion risk. Stacked bar plot shows that the proportion of mutations predicted to be at high reversion risk was 72%. **(J)** Application of the reversion-risk classification to clinical outcome in *BRCA2m* patients in the PAOLA-1 trial (*36*). Kaplan Meier plot depicts progression-free survival (PFS) in patients with *BRCA2* mutant high-grade serous ovarian cancer. Patients carrying pathogenic mutations classified as high reversion risk are shown in red, and those classified as low reversion risk in blue. *p* values were calculated using the log-rank test.

Focusing first on information that could be inferred by deletions in *BRCA1*, we noted that multiple *BRCA1* reverted patients had deletions that resulted in complete loss of the Serine-rich domain encoded by exon 10 (the largest exon, referred to as exon 11 in older literature; Fig. 4A, B Fig. 3D). Deletions in other previously defined functional BRCA1 domains were seen but these involved only part of the defined domain; for example, deletions in *BRCA1* that removed part of the RING finger, coiled-coil and BRCT domains were seen in the reversion cohort, but these domains were never completely deleted (Fig. 4A, B and described in detail later). For *BRCA2*, we observed numerous large deletions encompassing parts of exon 11, which encodes eight BRC repeats (Fig. 4C, D); these included the reversion seen in case KCL644_2 described above (Fig. 3A, B). Most BRCA2 domains annotated in the reversion cohort were disrupted, particularly the N-terminal DNA binding domain (N-DBD) and BRC repeats (Fig. 4C, D), but there was only one case where an annotated domain, the N-DBD of BRCA2, was completely deleted (KCL757_1, Fig. 4E, F).

In the second approach, we further defined the BRCA1 and BRCA2 regions that are rarely disrupted by reversion mutations (and thus likely to be more required for resistance to treatment) by calculating for each amino acid residue, the frequency at which it was observed to be disrupted (either by deletion, substitution, or being part of a frameshifted sequence) across all patients in the reversion cohort (the value C*j*, Fig. 5A-C). The cumulative sum of these frequencies along the protein was defined as the Total Reversion Impact (TRI). This residue-by-residue analysis allowed us to defined “cold regions” of BRCA1 or BRCA2 as those N- and C-terminal segments of each protein where amino acid residues were disrupted in less than 1% of the reverted patients (<1% TRI), and “hot regions” those regions of the proteins more frequently disrupted by reversions (Fig. 5A-C, for details see *Methods*).

For BRCA1, this analysis defined cold regions spanning residues 1-149 and 1608-1864, whilst the region 150-1607 was defined as hot (Fig. 5B). In addition, we noted that the entire coiled-coil domain of BRCA1, which interacts with PALB2 (described in detail later) and has recently been shown to be critical for PARPi resistance (*14, 15*), was largely preserved in reversions (Fig. 5B) despite being a common site for pathogenic mutations.

For BRCA2, two regions were defined as cold: residues 1-253 at the extreme N-terminus (“N-cold”) and the extreme C terminus (residues 2270-3418, “C-cold”, Fig. 5C). The relative rarity of disruptive reversions in the cold regions of BRCA1/2 was not obviously related to these regions lacking pathogenic mutations, with 35.6% of *BRCA1* and 34% of *BRCA2* pathogenic mutations expected to be in cold regions based on the incidence cohort (Fig. 5D). For *BRCA2*, pathogenic mutations in cold regions were underrepresented in the reversions cohort compared to their expected frequency in the incidence cohort (Fisher’s exact test, *p* = 3.5 x10^-6^, Fig. 5D). When reversions did occur in the BRCA2 “C-cold” region, these tended to be less disruptive and more conservative reversions, such as substitutions or true reversions (i.e. reversions to the wild type amino acid sequence), when compared to those found in the “Hot” region (Fisher’s exact test *p* = 4.4 x10^-11^ for C-cold *vs.* Hot, Fig. 5E). Furthermore, when more disruptive reversions in *BRCA2* did occur (e.g. deletions), those emerging from “cold” region pathogenic mutations had smaller deletion sizes compared with those in the “hot” region (Wilcoxon rank-sum tests *p* = 2.6 x 10^-2^ for N-cold and *p* = 1.5 x 10^-4^ for C-cold, Fig. 5F). An example of such a small, conservative deletion reversion in a *BRCA2* cold region is provided in Supplementary Figure S4, where a reversion in *BRCA2* removes one turn of an alpha helix, without disrupting the predicted protein folding or domain architecture.

As for *BRCA2*, pathogenic mutations in *BRCA1* cold regions were underrepresented in the reversion cohort *vs.* their frequency in the incidence cohort (Fisher’s exact test, *p* = 2.5 x10^-2^, Fig. 5G). When reversions did occur in N- or C-cold regions of BRCA1, these tended to be more conservative events (Fisher’s exact tests *p* = 1.1 x10^-3^ for N-cold and *p* = 4.3 x10^-3^ for C-cold, Fig. 5H) and when deletions, had shorter deletion lengths (Wilcoxon rank-sum test *p* = 1.6 x 10^-2^ for N-cold and *p* = 2.4 x 10^-5^ for C-cold, Fig. 5I).

These observations could imply that pathogenic mutations in hot regions have a heightened capacity to revert as they are able to revert via a wide range of mutations including those that have a highly disruptive effect on the local amino acid sequence; conversely there might be more constraints over pathogenic mutations in cold regions, that can only form functional proteins when reverted via more conservative reversion types, such as substitutions, true reversions or small deletions, when compared to those found elsewhere in the protein.

### Reversion cold spots define a requirement for PALB2 interactions in treatment resistance

Our above analysis also suggested that the extreme BRCA2 N terminal region that encompasses the PALB2 binding domain is “cold” and rarely disrupted by reversion, whereas the adjacent N-terminal DNA binding domain (N-DBD) was frequently disrupted by reversions or even completed deleted (Fig. 4C-F, Fig. 5C), suggesting N-DBD and the PALB2 binding domain might make different contributions to resistance. To functionally assess this possibility, we generated new cell line models of BRCA2 deficiency and introduced into these cDNA constructs expressing either wild type BRCA2 or BRCA2 forms lacking sections of the BRCA2 N-terminus informed by our analysis of the reversion cohort.

To facilitate the generation of *BRCA2* defective cells, we: (i) removed one copy of the *BRCA2* gene from diploid *TP53* defective RPE1-hTERT cells (haploidization, generating *BRCA2*^WT/–^cells); and then (ii) mutagenized the remaining wild type *BRCA2* allele using a CRISPR-Cas9 crRNA targeting a site in BRCA2 exon 11. This generated a *BRCA2* defective clone, *BRCA2*^c.4702insA/–^ that carried both a frame shift mutation, *BRCA2* ^c.4702insA^, and a *BRCA2* deletion.

*BRCA2* ^c.4702insA/–^ cells were PARPi sensitive and defective in their ability to form nuclear RAD51 foci in response to PARPi exposure (Fig. 6A-C, Figure S5) whilst re-introducing wild type *BRCA2* cDNA into these restored both PARPi resistance and RAD51 nuclear foci (Fig. 6D-H). PARPi resistance and RAD51 foci were also restored in *BRCA2* ^c.4702insA/–^ cells expressing a cDNA construct expressing the complete C terminus and five BRC repeats fused to a N terminus encompassing both PALB2 binding and N-DBD (construct ΔN1) as did a construct, ΔN2, which still encoded the N-terminal PALB2 binding domain, five BRC repeats and C terminus, but which lacked the N-DBD (Fig. 6D-G), an observation consistent with prior observations suggesting the N-DBD is not required for homologous recombination (*16*). Conversely, a construct ΔN3, which lacked both the N-DBD and the PALB2 binding domain did not restore PARPi resistance in *BRCA2* ^c.4702insA/–^ cells (Fig. 6D, E). Taken together, these data support the idea that the relative paucity of reversions in BRCA2 residues 1-254 when compared to those C-terminal to this “cold” region could be explained by the PALB2 interaction and N-DBD domains making different contributions to the PARPi resistance phenotype, with the N-DBD being non-essential for PARPi resistance, when compared to the PALB2 interaction domain.

Although rarer than *BRCA1/2* mutations, loss of function *PALB2* mutations also cause PARPi sensitivity (*17*). *PALB2* reversion mutations are also associated with PARPi resistance (*18, 19*). We therefore collated annotated and analyzed 33 *PALB2* reversion mutations from seven patients with treatment resistance (*19–25*) (Data S10, S11), reasoning that these might allow us to better understand the importance of the BRCA2/PALB2/BRCA1 interaction to PARPi resistance (Fig. S6A).

As for BRCA1 and BRCA2, most PALB2 reversions were deletions which allowed us to identify regions of the protein likely superfluous for PARPi resistance (Fig. S6B). Whilst several *PALB2* reversions deleted coding sequence for the central region of the PALB2 protein, the wild type coding sequence of the PALB2 C-terminal region that interacts with the BRCA2 N-terminus was always retained in reverted alleles (Fig. S6B). This was consistent with our prior analysis of BRCA2 reversions that indicated that the BRCA2/PALB2 interaction is critical for PARPi resistance.

PALB2 reversions also did not alter the amino acid sequence of the PALB2 N terminus that interacts with the BRCA1 coiled coil region (Fig. S6B), again consistent with our residue-by-residue analysis of BRCA1 reversions, which suggested that parts of the BRCA1 coiled coil region are rarely altered by reversions (Fig. 5B). This observation led us to examine in detail the few BRCA1 reversions that did alter the primary sequence of the BRCA1 coiled coil domain (Fig. S6C).

The BRCA1/PALB2 interaction involves two alpha helices which are orientated in an antiparallel fashion, one encompassed within the BRCA1 coiled coil domain (amino acids 1395-1422), the other formed by amino acids 11-63 of PALB2 (Fig. S6D). In totality, 12 BRCA1 reversions altered the BRCA1 coiled coil domain primary sequence (Fig. 4B, Fig. S6C); of these, seven were deletions affecting only the first two amino acids of the domain (encoded by exon 10) and were thus predicted to not alter the BRCA1 alpha helix encoded in exon 11 (Fig. S6C). Each of the other BRCA1 reversions that altered the coiled coil domain either: (i) modified a disordered region of BRCA1 which was N-terminal to the alpha helix, without disrupting the predicted folding of the alpha helix itself (Fig. S6E-G); or (ii) changed the primary sequence of the alpha helix without obviously perturbing its predicted tertiary structure (Fig. S6H,I). As we describe later, CRISPR-Cas9 screens that deleted large portions of *BRCA1* but reverted the gene always produced PARPi resistant cells that retained the BRCA1 coiled coil alpha helix coding sequence. Taken together, with the BRCA2 and PALB2 reversion data, this implied that one constraint on BRCA1, BRCA2 and PALB2 reversions is that they retain the coding sequences that allow BRCA1/PALB2 and BRCA2/PALB2 interactions to occur.

### PARPi resistance can be mediated by BRCA2 forms that lack exon 11 encoded BRC repeats

Most BRCA2 reversions (383/606 reversion events, 63%) occurred in the central portion of the coding region, specifically in exon 11, which encodes the 8 BRC repeats (Fig. 4C, D, Fig. 5C, Fig. 7A) which functionally are involved in binding to the recombinase RAD51 in its monomeric state (*26, 27*). In patients with *BRCA2* exon 11 pathogenic mutations, we also identified reversions that deleted the exon 11 splice donor site (e.g. case Simmons 12, Fig. 7B); this splice donor reversion could potentially lead to loss of the entire exon 11 sequence in mature mRNA, for example by in-frame splicing from exon 10-12 or 10-13 (Fig. 7B). Given this, we set out to assess whether resistance to therapy could be mediated by reversion events that remove all BRCA2 exon 11 encoded BRC repeats. Functionally, such a scenario might be possible: although the BRCA2 exon 11 encoded BRC repeats are critical for RAD51 filament formation, recent data suggest that the C-terminus of BRCA2, which stabilizes RAD51 nucleoprotein filaments (*28*), is more important for both homologous recombination and PARPi resistance (*29*).

To characterize whether whole exon deletion (of exon 11 or indeed any other exon) is a viable route to cellular PARPi resistance, we carried out an unbiased CRISPR-Cas9 mutagenesis screen using a library of 2910 dual-targeting sgRNA vectors where each vector was designed to cut one pair of *BRCA2* introns (e.g. one sgRNA pair designed to cut *BRCA2* intron 2 and intron 3, Fig. 7C). An average of three guides per intron were used, and the library also included 60 control vectors that encoded either non-targeting guides or those that were known to cause lethality. This sgRNA library was used to mutagenize the *BRCA2* defective *BRCA2* ^c.4702insA/–^ cells and also a second PARPi sensitive clone with a BRCA2 exon 11 frameshift mutation that was derived from mutagenesis of BRCA2 ^WT/–^ cells, *BRCA2* ^c.4701_4729del/–^ (Fig. S5). Cells were then exposed to one of two different clinically used PARPi, olaparib or talazoparib for two weeks, after which sgRNA pairs present in the surviving, PARPi resistant, cell population were identified by deep sequencing (Fig. 7D).

Both *BRCA2* ^c.4702insA/–^ and *BRCA2* ^c.4701_4729del/–^ screens showed that dual sgRNA pairs designed to delete both exons 11 and 12 (i.e. guide pairs that simultaneously cut in intron 10 and 12) were consistently enriched in the PARPi surviving population, whether olaparib (Fig. 7E) or talazoparib (Fig. 7F) exposure was used, consistent with the idea that BRCA2 exon 11 might be superfluous for PARPi resistance. We therefore validated the most enriched dual guide construct by transfecting *BRCA2* ^c.4702insA/–^ cells with this construct alone, followed by selection in the novel PARP1-selective inhibitor saruparib (*30*) (Fig. 7G). In the resultant PARPi resistant cells, termed *BRCA2* ^ΔEx11-12/–^ cells (Fig. 7H), new *BRCA2* deletion break points within introns 10 and 12 were detected by DNA sequencing, consistent with the loss of exons 11 and 12. We also confirmed expression of a BRCA2 protein from this new deleted allele using an antibody that detected a C-terminal epitope; as predicted from DNA sequencing, *BRCA2* ^ΔEx11-12/–^ cells expressed a BRCA2 protein of 198 kDa that possessed the native BRCA2 C-terminus (Figure 7I).

*BRCA2* ^ΔEx11-12/–^ cells possessed a similar level of PARPi resistance as locally haploid BRCA2 ^WT/–^cells that retained a BRCA2 wild type allele (Fig. 7J). *BRCA2* ^ΔEx11-12/–^ cells also had increased levels of RAD51 foci than in BRCA2 defective *BRCA2* ^c.4702insA/–^ cells (Fig. S7A-E). We also demonstrated that BRCA2 was responsible for both PARPi resistance and RAD51 foci in *BRCA2* ^ΔEx11-12/–^ cells by silencing of BRCA2, which reversed both phenotypes (Fig. S7F-H). Consistent with the idea that BRCA2 proteins lacking all BRC repeats could mediate resistance, we also designed a cDNA construct that lacked all exon 11 BRC repeats (Fig. S8A, ΔBRC1-8). This construct partially rescued PARPi resistance (Fig. S8B) and RAD51 foci formation (Fig. S8C, D) in *BRCA2*^c.4702insA/–^ cells, albeit to a degree less than in cells transfected with a wild type BRCA2 expression construct or cells expressing a BRCA2 cDNA encompassing at least one BRC domain (Fig. S8A, E). Importantly, we do not hypothesize that the predicted minimal forms of BRCA2 such as those encoded by exon 11-12 deleted alleles are required for all functions of the protein but perhaps only those relevant to resistance to PARPi. For example, for BRCA2, the full complement of BRC-repeats might not be required for PARPi resistance but might be required for BRCA2 to respond to other forms of DNA damage or required for other functions of the protein, such as those critical to meiosis.

### The BRCA1 coiled-coil domain is retained in BRCA1 reversions

To investigate which domains of BRCA1 might be functionally critical for PARPi resistance, we applied a similar dual CRISPR screen approach to *BRCA1* as used above for *BRCA2*. Here we used SUM149 cells, a triple negative breast tumor cell line which has a pathogenic *BRCA1* mutation in exon 10. These cells were mutagenized with a bespoke dual guide sgRNA library encoding 2,080 pairs of sgRNAs, with each pair designed to cut one pair of *BRCA1* introns. Cells were then exposed PARPi for two weeks as before, to identify large BRCA1 deletion events that caused PARPi resistance.

In this case, the most profound effects seen in the PARPi SUM149 population involved dual sgRNA vectors that encoded guides cutting either side of exon 10, as well as potential deletions of exons 8 to 10 and exons 6 to 10 (Fig. S9). These events encompass most of the BRCA1 “hot” region, thus functionally supporting the idea that this is not essential for PARPi resistance. Expression of BRCA1 variants lacking all, or the majority of exon 10 (referred to as exon 11 in historical transcript annotation) such as the Δ11q splice variant, have previously been shown to confer PARPi resistance (*31*). Notably, deletions in our BRCA1 dual CRISPR screen did not extend beyond the coiled-coil domain (see earlier and Fig. S9C), which in our prior amino acid by amino acid analysis was shown to be in a cold region of the protein largely preserved in reversions (Fig. 5B), and is known to be critical for PARPi resistance (*14, 15*).

### Design of a prototype predictor of *BRCA2* reversion risk

One of our aims in studying *BRCA1/2* reversions was to identify the rules that govern the probability of a pathogenic *BRCA1/2* mutation to revert. Since the presence of reversion mutations is associated with shorter time to disease progression (*32*), being able to predict such a property could serve as a biomarker of patient outcome, by either estimating how much benefit each patient might achieve from a PARPi, whether enhanced surveillance for the emergence of reversions using ctDNA sampling might be warranted, or whether a reverted and potentially fitter and thus more lethal tumor subclone might rapidly emerge. As a simple example that the propensity to revert could have clinical value, we note that ovarian cancer patients with large genomic deletions in *BRCA1/2* (i.e. *BRCA1/2* alterations that are unable to revert) tend to exhibit long-term responses to PARPi (*33*), as do missense pathogenic mutations in RAD51C, which also controls homologous recombination (*34*). Furthermore, ovarian cancer patients with pathogenic mutations in the BRCA2 C-terminus (which are significantly underrepresented in the reversions cohort despite being well represented in the incidence cohort – see earlier) also have better outcomes when treated with maintenance PARPi (*35*). Based on the analysis of our expanded reversions cohort and the observations described above, we formulated a series of heuristics or rules that allowed us to model reversion risk for a given pathogenic *BRCA1* or *BRCA2* mutation (Fig. 8A). These heuristics included: (i) whether a pathogenic mutation has previously been observed to revert (i.e. was the pathogenic mutation in our reversion cohort); (ii) the type of pathogenic mutation present (e.g. missense and splice mutations appear less likely to revert since these are underrepresented in the reversions cohort compared with the incidence population, Fig. 2A); (iii) whether the pathogenic mutation is located in “hot” or “cold” regions based on our residue-by-residue analysis described in Fig. 5. We encapsulated all these heuristics into an algorithm that classifies pathogenic mutations into high or low reversion risk (Fig. 8A). Applying this to our incidence cohort classified approximately 72% of pathogenic mutations at high risk of reversion (Fig. 8B, Data S12). We then assessed whether this classification of reversion risk: (i) correlated with reversion frequency in two different clinical datasets; and (ii) correlated with response to drugs that are used to target the DNA repair defect in *BRCA1/2* mutant cancer, as measured by progression free survival in patients receiving platinum salts and PARP inhibitors in two clinical studies. Although most reports identifying *BRCA1/2* reversions only describe small numbers of patients and focus on patients that did revert, these studies, BTBC (*5*) and the ARIEL2 trial (*11*), are the largest reported studies, in terms of patient numbers, where reversion mutations have been analyzed but resistant patients without reversions are also reported. BTBC was a biomarker study where reversions were studied in metastatic breast cancer patients with germline *BRCAm* who received either the PARPi olaparib and/or platinum salts as part of their standard of care treatment (*5*) whereas ARIEL2 was a phase 2 clinical trial where *BRCAm* ovarian cancer patients received platinum salt treatment followed by the PARPi rucaparib: in ARIEL2 patients were retrospectively assessed for reversion mutations (*11*). After omitting data from either BTBC or ARIEL2 patients in step 1 of the predictor (prior observation of reversion), we classified all *BRCA1/2* pathogenic mutations into high or low reversion risk. Similar to the incidence cohort, the proportion of patients at risk of reversion observed in the BTBC and ARIEL2 cohorts were 76% (BTBC, Data S13) and 73% (ARIEL2 Data S14) (Fig. 8C, D). We then assessed whether high or low risk of reversion classifications correlated with reversion observation in both studies (Data S13, S14). In both BTBC and ARIEL2 patients, those classified as having pathogenic *BRCAm* at high risk of reversion exhibited a higher frequency of reversion than those classified as having pathogenic mutations at low risk of reversion (Fig. 8E-G, Data S15).

Although BTBC is not a clinical trial and thus we were unable to correlate reversion risk with clinical outcome on HRD-targeting therapies, some insight can be gained from ARIEL2, where the platinum free interval (PFI, Data S14) for each patient, a measure of response to the last line of platinum therapy prior to entering the ARIEL2 trial, has previously been reported (*11*). We found that *BRCA2m* patients in ARIEL2 with pathogenic mutations at low risk of developing reversion showed a longer PFI, than those with *BRCA2* mutations classified as high risk of reversion (Fig. 8H; *p* = 0.039), consistent with the hypothesis that for *BRCA2m* patients, the response to HRD-targeting therapy might be linked to risk of reversion. A similar correlation was not observed for *BRCA1* carriers (Figure S10A), possibly because more mechanistic routes to drug resistance other than *BRCA*m reversion exist for *BRCA1* mutant cancers than for *BRCA2* mutant cancers (see *Discussion*). The only other clinical trial where HRD-targeting therapy has been assessed and where the pathogenic *BRCA*m of each patient has been disclosed is the PAOLA-1/ENGOT-ov25 phase 3 clinical trial (PAOLA-1) (*36*), within which 233 patients with *BRCA*m ovarian cancer who responded to platinum-taxane chemotherapy in combination with the VEGF-targeting antibody bevacizumab were next treated with either bevacizumab or bevacizumab combined with the PARPi olaparib (*35*). Although patients were not directly assessed for reversion mutations in this trial, patients with *BRCA2* pathogenic mutations classified as low risk of reversion had better progression-free survival (PFS, Fig. 8J, K, Data S16), similar to the PFI relationship observed in ARIEL2 above. However, as for ARIEL2, there was no significant difference in PFS when the classifier was applied in patients with *BRCA1* mutations (Fig. S10B). Together, these findings suggest that for *BRCA2*, the risk of reversion might correlate with both the presence of reversions and treatment outcome.

## Discussion

In this analysis, we describe 384 reported cases of clinical reversion mutations in *BRCA*m cancer patients with resistance to HRD targeting therapy. A large number of reversion mutations result in disruption (by deletions, substitutions, or frameshifted sequences) of BRCA1/2 protein coding sequence. Analyzing the regions that are recurrently disrupted or preserved by reversions gives clues to the rules governing the emergence of reversion mutations from a pathogenic *BRCA1/2* mutation, and the functional domains of BRCA1/2 that are dispensable for promoting resistance to PARPi. It is particularly striking that the large central exon of each gene is broadly dispensable in terms of PARPi resistance.

The conclusions of our analysis have some key similarities and differences to studies describing the requirements for *BRCA2* in cellular homologous recombination (HR), one of the DNA repair processes ascribed to *BRCA1* and *BRCA2*. For example, by combining analysis of our reversion dataset with functional analysis, we show that exon 11 of BRCA2 is largely dispensable for PARPi resistance but that the extreme N and C termini of the protein are not. These observations, made from the analysis of cancer patients with resistance to treatments that target defective homologous recombination, concur with data generated in cell lines expressing different *BRCA2* mutants and biochemical analysis of *BRCA2* that suggests that the BRCA2 C-terminus is critical for homologous recombination and PARPi resistance (*28, 29, 37*), whilst RAD51 loading, the canonical function of the BRC repeats encoded by exon 11 can also be mediated via BRCA2-independent means. In other work, at least two BRC domains, plus the flanking N and C terminal domains of *BRCA2*, have been shown to be required for maximal repair of an enzymatically generated double strand break in an HR reporter gene, DR-GFP (*16*). Rather than expecting these observations to be contradictory, what is perhaps more likely is that these reflect differing requirements for BRCA2 domain to repair a chromosomal double strand DNA break that is not necessarily associated with replication forks (as is measured using DR-GFP) with the DNA lesion or lesions caused by PARP inhibitors. In a similar vein, the functions of BRCA1/2 required for resistance to therapy in a patient may not always be the same as the functions required for tumor suppression, or for other cellular functions. For example, in mice, complete loss of exon 11 encoded BRC repeats causes embryonic lethality (*38, 39*) and distant orthologues of BRCA2 have varying numbers of BRC repeat domains, but always encode at least one (*40, 41*) suggesting that there are some evolutionary conserved functions of BRCA2 in development that have a BRC requirement; PARPi resistance (which presumably is an evolutionarily new biological phenotype) might not have the same functional requirements. As such, our work defines the minimal BRCA1/2 requirements for PARPi resistance rather than the minimal requirements for specific forms of DNA repair or tumor suppression *per se*.

Interestingly reversion risk was associated with outcome for *BRCA2* mutant patients, but not for those with *BRCA1* mutations. In considering why this might be, we note that in addition to *BRCA1* reversion, a number of additional non-reversion-based mechanisms of PARPi resistance have been identified in *BRCA1* mutant models or patients, that might not be expected to cause PARPi resistance in *BRCA2* mutant cancers, such as loss of TP53BP1 or the Shieldin complex (*42, 43*) or alternative splicing of *BRCA1* (*44–46*). If additional mechanisms of PARPi resistance other than *BRCA*m reversion play a greater role in *BRCA1* as opposed to *BRCA2* mutant cancers, the relative contribution of reversion to the overall clinical outcome of patients might be greater for *BRCA2* than for *BRCA1*. As such, the asymmetry between *BRCA1* and *BRCA2* in terms of whether reversion risk is associated with outcome could reflect fundamental differences in the landscape of resistance mechanisms available to tumours with mutations in each gene.

Previous studies have suggested that pathogenic mutations in *BRCA1/2* and other HR genes have constraints on their ability to revert, such as missense mutations which can most likely only revert via a compensating missense mutations that restore the wild type amino acid or via mutations that have very similar biochemical properties (*34, 47, 48*). Large genomic deletions removing extensive coding sequence have also been associated with good outcomes in PARPi trials, which could be due to their inability to revert (*33*). We sought here to develop a more generalized predictor of reversion risk using rules derived from our analysis of clinically observed *BRCA1/2* reversions. One implication of identifying cancer patients with *BRCA1/2* pathogenic mutations more likely to revert is that this might eventually identify the cohort of patients who might benefit from more intensive screening for reversions whilst on PARPi or platinum salt treatment (for example, by routine ctDNA sequencing) as an approach to identify the early emergence of resistant disease. In the United States alone, breast, prostate, pancreatic, and ovarian cancers together are estimated to account for over 719,000 cases per annum (SEER, National Cancer Institute). Among these, approximately 10% are expected to harbor *BRCA1/2* mutations (about 71,900 cases), and based on our analysis, around 72% of these patients (about 52,000 cases per annum) are predicted to be at high risk of developing reversion mutations. These figures support the need for close molecular monitoring to anticipate treatment resistance and thus the further assessment of prototypical reversion risk predictor such as that described here.

Finally, our data indicate that very minimal forms of BRCA1, and especially BRCA2, appear able to cause drug resistance. Over the course of evolution, the size of the BRCA1 and BRCA2 (and most other proteins) has increased, most likely as doing so allows proteins to facilitate additional functions or to be more amenable to fine tuning of existing functions. Our data indicate that in tumor cells with PARPi or platinum resistance, BRCA1 and BRCA2 start to resemble forms seen in lower organisms. This scenario seems reminiscent of *adaptive reversion,* where tumor cells revert to a previous, ancestral, state that provides a survival advantage. It seems reasonable to think that other forms of drug resistance in cancer might also operate via adaptive reversion and that studying this as a cause of drug resistance could provide insight into drug resistance and the evolution of cancer more generally.

## Acknowledgments

We thank S. Jackson (University of Cambridge, UK) for provision of RPE1 TP53^−/–^ cells. We thank S. Nik-Zainal and D. Black (University of Cambridge, UK) for critical review of the work and useful discussions.

## Funding

Programme Grant from Breast Cancer Now as part of Programme Funding to the Breast Cancer Now Toby Robins Research Centre to CJL, SP and ANJT.

Programme Grant from Cancer Research UK to CJL, SP and ANJT.

Marie Sklodowska-Curie Postdoctoral Fellowship funded by UK Research and Innovation (UKRI) under the UK government’s Horizon Europe funding guarantee grant EP/Y010361/1 (LMP)

This work represents independent research supported by the National Institute for Health Research (NIHR) Biomedical Research Centre at The Royal Marsden NHS Foundation Trust and the Institute of Cancer Research, London. The views expressed are those of the author(s) and not necessarily those of the NIHR or the Department of Health and Social Care.

## Author contributions

Conceptualization: CJL, SP, ANJT

Methodology: all authors Investigation: all authors

Funding acquisition: LMP, CJL, SP, ANJT

Supervision: CJL, SP, ANJT

Writing – original draft: CJL, SP, ANJT

Writing – review & editing: all authors

## Competing interests

CJL makes the following disclosures: receives and/or has received research funding from: AstraZeneca, Merck KGaA, Artios, Neophore, FoRx. Received consultancy, SAB membership or honoraria payments from: FoRx, Syncona, Sun Pharma, Gerson Lehrman Group, Merck KGaA, Vertex, AstraZeneca, Tango Therapeutics, 3rd Rock, Ono Pharma, Artios, Abingworth, Tesselate, Dark Blue Therapeutics, Pontifax, Astex, Neophore, Glaxo Smith Kline, Dawn Bioventures, Blacksmith Medicines, FoRx, Ariceum. Has stock in: Tango, Ovibio, Hysplex, Tesselate, Ariceum. C.J.L. is also a named inventor on patents describing the use of DNA repair inhibitors and stands to gain from their development and use as part of the ICR “Rewards for Discoverers” scheme and also reports benefits from this scheme associated with patents for PARP inhibitors paid into CJL’s personal account and research accounts at the Institute of Cancer Research.

ANJT reports personal honoraria for advisory boards from AstraZeneca, ForEx Therapeutics, Nauge Therapeutics and Artera, and honoraria for advisory boards to either The Institute of Cancer Research or King’s College research accounts from AstraZeneca Page Therapeutics, Ellipses, Tango Therapeutics, Guardant Health, Merck KGaA, as well as speaker fees from AstraZeneca and Permanyer Mexico. AT reports financial support for research from AstraZeneca, Myriad Genetics, Merck KGaA and compensation for travel expenses from AstraZeneca. AT reports benefits from ICR’s Rewards for Discoverers Scheme associated with patents for PARP inhibitors in BRCA1/2 associated cancers, paid into research accounts at The Institute of Cancer Research and to AT’s personal account.

SJP makes the following disclosures: SJP is a named inventor on patents describing the use of DNA repair inhibitors and treatment of cancers with reversion mutations and stands to gain from their development and use as part of the ICR “Rewards for Discoverers” scheme. SAB membership: VUS Genetics.

IR-C makes the following disclosures: Grants: Bristol Myers Squibb, MSD Oncology, Roche/Genentech. Consulting fees: AbbVie, Agenus, AstraZeneca, Antares, Blueprint Medicines, Bristol Myers Squibb, Clovis Oncology, Corcept, Daiichi Sankyo, Deciphera, Eisai, Genmab, GlaxoSmithKline, Immunocore, Immunogen, MacroGenics, Lilly, Mersana, MSD Oncology, Novartis, Novocure, Ose Immunotherapeutics, PharmaMar, Roche, Pfizer, Seagen, Scorpion, Sutro Biopharma, Tesaro. Honoraria: AbbVie, Advaxis, Agenus, Amgen, AstraZeneca, Bristol Myers Squibb, Clovis Oncology, Daiichi Sankyo, Deciphera, Genmab, GlaxoSmithKline, Immunocore, Immunogen, MacroGenics, Mersana, MSD Oncology, Novartis, OxOnc, Pfizer, PharmaMar, PMV Pharma, Roche, Seagen, Sutro Biopharma, Tesaro. Meeting/travel support: Abbvie, AstraZeneca, Bristol-Myers Squibb, Clovis Oncology, GlaxoSmithKline, Esai, PharmaMar, Roche. Other: GINECO, Belgium Health Authorities, French National Cancer Institute (INCa), German Health Authorities, Italian Health Authorities (uncompensated) E P-L makes the following disclosures: employee of ARCAGY-GINECO.

All other authors declare that they have no competing interests.

## Data and materials availability

All data, code, and materials used in the analysis are either provided as part of the supplementary material, available via https://software.icr.ac.uk/app/reversions-web, or available upon reasonable request to the corresponding authors. Access to data from PAOLA-1 (*36*), is made via request to the PAOLA-1 translational research committee.

## Materials and Methods

### Compilation of the reversion cohort

The reversions cohort was curated via systematic analysis of peer reviewed literature (PubMed search, cutoff December 2024, Fig. S1, Data S1) that described *BRCA1/2* reversion mutations and corresponding *BRCA1/2* pathogenic mutations in breast, ovarian, prostate or pancreatic cancer patients. For each event, available clinical and molecular metadata were retrieved, including tumour histology, treatment used, sequencing technology, material analysed (e.g., circulating tumour DNA, tumour biopsy, ascites) when reported. A dedicated curation step was implemented prior to downstream analyses. For each event, the allele containing both the pathogenic mutation and the corresponding reversion was systematically transcribed at the transcript level using an automated pipeline to assess whether the coding sequence was restored. Events that did not restore the reading frame were flagged and excluded from downstream analyses. All variants were standardized to HGVS nomenclature using a single reference transcript per gene. Transcript-level consequence annotations and genomic coordinates (hg38) were obtained using Ensembl Variant Effect Predictor (VEP release 114, May 2025 (*49*)) and ANNOVAR (*50*). This curated and standardized database formed the basis for subsequent systematic analyses of reversion patterns. Full details of the reversions cohort are described in a publicly accessible database, https://software.icr.ac.uk/app/reversions-web.

### Compilation of the incidence cohort

We built an incidence dataset (Data S7) of 1018 patients to serve as a reference population for comparison with the reversion cohort. This dataset included patients with *BRCAm*-associated cancers who were considered for platinum-based or PARPi therapy. For the incidence cohort, the pathogenic *BRCA*m for each patient was retrieved. Clinical information, including tumour type was also collated. All variants were standardized to HGVS nomenclature using the same reference transcripts applied to the reversion cohort to ensure consistency across analyses.

### Characterization of reversion types

Reversion events were initially annotated using Variant Effect Predictor (VEP; Ensembl release 114, May 2025 (*49*)) to obtain standardized transcript-level consequence annotations. As reversion mutations do not constitute a predefined functional category, events were subsequently manually curated and classified according to their predicted mechanism of functional restoration. Reversions were categorized as: (i) inter-exon deletions involving both exonic and intronic sequences; (ii) secondary frameshift mutations restoring the open reading frame; (iii) intra-exon deletions removing in-frame coding sequence encompassing the original pathogenic mutation; (iv) secondary substitutions, defined as point mutations occurring within the same codon as the pathogenic variant and restoring the affected codon; (v) true reversions, i.e. reversion to the wild type sequence; (vi) splice reversion mutations generating alternative transcripts predicted to restore the reading frame (vii) other events not fitting these prior definitions.

### Deletion analysis

Clinically observed reversion events involving deletions, previously classified as inter-exon deletions, intra-exon deletions or secondary frameshift mutations, were mapped along the linear BRCA1 and BRCA2 protein sequences according to their corresponding amino acid coordinates. For secondary frameshift mutations, the region spanning the original pathogenic frameshift mutation and the secondary frameshift mutation that restored the reading frame was defined as the frameshift-altered coding region.

### Residue-by-residue analysis

To quantify how frequently each amino acid is disrupted by clinical reversion events, we created a binary matrix Mij, where i represents the revertant cases (from 1 to n) and j the amino acid position in BRCA2 (from 1 to 3418). The matrix was populated such that, per patient (i), each residue (j) disrupted by reversion (i.e., deleted, substituted, or part of a frameshifted sequence) was marked with 1, while residues not altered by a reversion in each patient were marked with 0.

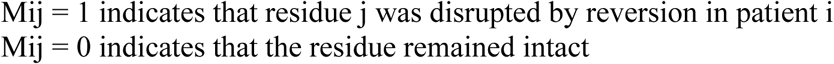

To avoid overcounting in patients with multiple reversions events, the matrix was normalized per patient by collapsing all reversions into a series of single patient-centric profiles, so that each disrupted amino acid residue was counted only once per patient. The cumulative sum per amino acid (Cj) reflects how frequently each residue is disrupted per patient. The same strategy was used to generate the residue-by-residue analysis in BRCA1.

### Hot/Cold regions definition

From the residue-by-residue analysis, we calculated the cumulative number of amino acid positions disrupted by reversion events across all patients. This total count was defined as the Total Reversion Impact (TRI).

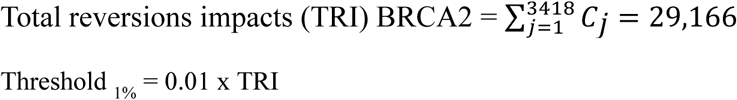

Cold regions were defined as the N- and C-terminal segments of BRCA2 protein where the cumulative number of disrupted residues, starting from each extreme, reached the threshold defined as 1% of the TRI. The hot region was defined as the central segment encoded between the N-Cold and C-Cold boundaries.

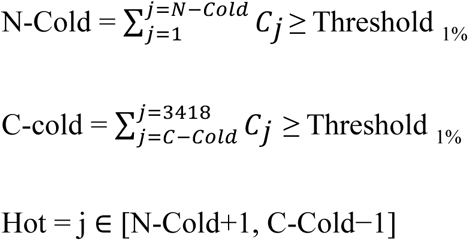

The same approach was applied to BRCA1 to define hot and cold regions.

### Microhomology analysis

Deletion-mediated reversions (excluding indels) were selected for microhomology analysis if deletion lengths were >1 bp in length and where the precise breakpoint of the deletion was identified. Events where the reversion start position overlapped with the pathogenic mutation breakpoint were excluded for the analysis. In-house R scripts were used to systematically determine the presence of microhomology (μH) flanking the deletion junctions, as well as subsequent data exploration.

### Generation of BRCA2 haploid cells and cells with BRCA2 exon 11 mutations

hTERT RPE-1 *TP53* ^−/−^ cells (the kind gift of S. Jackson, University of Cambridge) and derivatives were maintained in DMEM/F-12 supplemented with 10% (v/v) heat-inactivated fetal bovine serum (FBS) and 1% (v/v) penicillin–streptomycin. SUM149 cells were cultured in Ham’s F-12 medium containing 5% (v/v) FBS, 5 µg/mL insulin, and 1 µg/mL hydrocortisone. All cells were cultured at 37 °C in a humidified incubator with 5% CO₂ and dissociated with TrypLE Express for routine passaging.

To generate RPE-1 *TP53 ^−/−^* cells carrying a single *BRCA2* allele, short guide RNAs targeting genomic regions flanking the *BRCA2* locus (BRCA2 gRNA/1 and BRCA2 gRNA/1, Data S17) were complexed with recombinant Cas9 protein and tracrRNA (Thermo Fisher Scientific) and delivered into cells using CRISPRMAX Lipofectamine (Invitrogen) according to the manufacturer’s instructions. Seventy-two hours post-transfection, single cells were isolated and expanded clonally.

Genomic DNA was extracted from expanded clones using QuickExtract DNA Extraction Solution (Biosearch Technologies). Deletion of *BRCA2* was confirmed by PCR and Sanger sequencing using primers flanking the spacer regions (BRCA2 F1 and BRCA2 R1, Data S17) as well as using PCR/sequencing primers spanning a heterozygous BRCA2 SNP (BRCA2 SNP F and BRCA2 SNP R, Data S17; Supplementary Fig. 5A–B). Relative copy number was quantified using droplet digital PCR (ddPCR) with probes targeting BRCA2 and reference genes RNase P and FOLR1 located on separate chromosomes. Droplet generation and reading were performed using the QX200 system (Bio-Rad) according to the manufacturer’s instructions (Supplementary Fig. 5C).

*BRCA2* exon 11 mutations were introduced into *BRCA2* ^WT/−^ cells by transfecting these with a Cas9–crRNA ribonucleoprotein targeting exon 11 (BRCA2 gRNA/3, Data S17) using CRISPRMAX Lipofectamine. Single-cell clones were isolated and genotyped by PCR and Sanger sequencing using primers BRCA2 F2 and BRCA2 R2 (Data S17).

### cDNA transfection

Wild-type BRCA2 cDNA was cloned into a PiggyBac transposon vector carrying a neomycin resistance cassette. Mutant BRCA2 cDNAs carrying defined domain deletions were generated by site-directed mutagenesis. All constructs were validated by nanopore sequencing to confirm the expected deletions and exclude off-target variants. RPE-1 cells were co-transfected with PiggyBac constructs and the hyPBase transposase (*51*) using Lipofectamine 3000 (Thermo Fisher Scientific) according to the manufacturer’s instructions. Stable integrants were selected using 500 µg/mL geneticin.

### RAD51 foci quantification

Cells were exposed to 5 µM olaparib for 24 hours or received 10 Gy of ionizing radiation prior to fixation. Cells were fixed with 4 % (v/v) paraformaldehyde, permeabilized with 0.2 % (v/v) Triton X-100, and blocked with 1 % (w/v) bovine serum albumin (BSA) and 2 % (v/v) fetal bovine serum (FBS). Cells were incubated overnight at 4°C with rabbit anti-RAD51 (EPR4030(3); Abcam) and mouse anti-γH2A.X (JBW301; ThermoFisher Scientific). After washing, cells were incubated with Alexa Fluor 647–conjugated donkey anti-mouse (A-31573; ThermoFisher Scientific) and Alexa Fluor 555–conjugated donkey anti-mouse (A-31570; ThermoFisher Scientific) secondary antibodies for 1 hour at room temperature. Nuclei were counterstained with DAPI. Nuclei were counterstained with DAPI. Images were acquired on the Opera Phenix high-content imaging system. RAD51 foci–positive cells were defined as nuclei containing ≥5 RAD51 foci. Quantification was performed using Harmony (Revvity) software.

### PARPi dose response estimation

Cells were seeded in 24-well or 96-well plates at densities of 250 or 200 cells per well, respectively. After 24 h, cells were exposed to a range of drug concentrations. Media containing fresh drug were replenished every 3–4 days. After 6 or 10 days of drug exposure, cell viability was estimated using the CellTiter-Glo (Promega) luminescent assay. Luminescence values were normalized to DMSO controls to calculate surviving fractions. Dose–response curves were fitted using a four-parameter logistic model in GraphPad Prism 9.

### Dual CRISPR screens

A custom dual sgRNA lentiviral library targeting combinations of BRCA1 or BRCA2 introns, established synthetic lethal genes, single essential genes, or safe controls was constructed (three sgRNAs per intron or control gene; sequences in Data S 18). The library was cloned into a lentiviral vector carrying a puromycin resistance cassette (pKLV2_U6sgRNA_Puro2AmAG1_CmR-ccdB, modified from Addgene #67976). Cells were transduced at a multiplicity of infection of 0.3 (to prevent multiple integrants per cell) and selected with puromycin (2 µg/mL for RPE-1; 1 µg/mL for SUM149) for three days, after which cells were exposed to DMSO, olaparib (5 μM for RPE1 cells, and 2 μM for SUM149 cells), or talazoparib (50 nM for RPE1 cells, and 20 nM for SUM149 cells) for 14 days after which genomic DNA was isolated from the surviving population using the Puregene kit (Qiagen). sgRNA cassettes were PCR-amplified (using primers Exon Screen F and Exon Screen R, Data S 18) and subjected to next-generation sequencing to quantify reads per guide. Z-scores or guide read counts were calculated per guide and normalized to pre-treated screen samples to identify sgRNA pairs whose abundance significantly changed after PARPi exposure.

### Immunoblotting

Cells were lysed in RIPA buffer (ThermoFisher Scientific) supplemented with protease inhibitors (Roche). Protein concentration was measured using the Pierce BCA assay. Immunoblotting was performed using the Jess Simple Western system (ProteinSimple) with the following antibodies: anti-BRCA2 (D9S6V; Cell Signaling Technology), anti-vinculin (7F9; Santa Cruz Biotechnology), and anti-GAPDH (60004-1-Ig; Proteintech). Chemiluminescent or infrared detection modules (Bio-Techne) were used according to the manufacturer’s instructions.

### Reversion risk prediction

An algorithm was designed to classify pathogenic mutations into high or low reversion risk using heuristics directly derived from the reversion cohort, including clinically observed reversion events. First, mutations previously observed to undergo reversion and annotated in the reversion cohort were directly classified as high reversion risk. If no prior reversion had been reported, mutations were classified according to their type. Splice-site, missense mutations, and structural variants were directly classified as low reversion risk, as their prevalence in the reversion cohort is significantly lower compared with the population of *BRCA*-mutant patients treated with platinum or PARPi (incidence population). On the opposite, frameshift and stop-gain mutations were classified as high or low reversion risk depending on their location within the protein sequence. Mutations located in hot regions were classified as high risk, whereas those located in cold regions were classified as low risk.

### Analysis of BTBC, ARIEL2 and PAOLA data

Pathogenic mutations in patients from the incidence cohort, BTBC (*5*), ARIEL2 (*11*), and the PAOLA-1 trial (*36*) were classified as high or low reversion risk using the reversion risk classifier. For BTBC and ARIEL2, step 1 of the classifier (reversion history) was omitted for the purposes of this analysis. Three cases from PAOLA-1 trial were excluded due to ambiguous annotation. Predictive performance of the reversion risk classifier was evaluated in BTBC and ARIEL2, overall and in gene-stratified subsets (*BRCA1* and *BRCA2*). Predicted reversion likelihood (High vs Low) was compared with observed reversion status. Performance was quantified using F1-score, sensitivity, and specificity. Metrics were calculated independently for each dataset in R (version 4.4.1) using the PerfMeas package. The association between predicted reversion risk (High or Low) and progression-free survival was evaluated in the ARIEL2 and PAOLA-1 trials, including gene-stratified analyses (*BRCA1* and *BRCA2*). Kaplan-Meier curves were compared using log-rank tests. Analyses were performed in R using the survival and survminer packages.

## Supplementary Figures

**Fig. S1.**
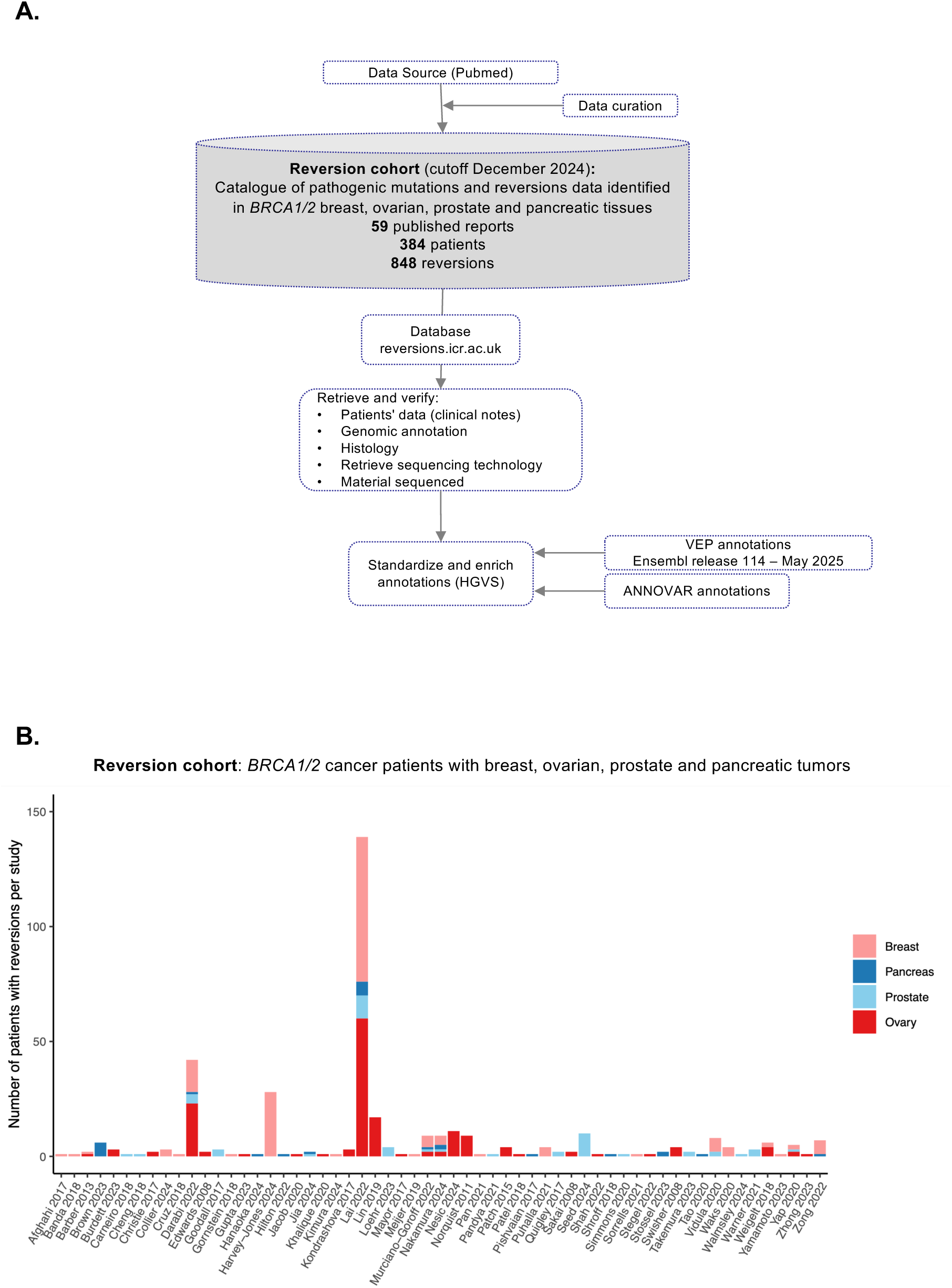
Pipeline for reversion mutation curation and analysis. **(A)** Flowchart illustrating the pipeline used to compile, annotate, and analyze reversion mutations. Data from 59 publications were curated and incorporated into a comprehensive catalogue of 848 BRCA1/2 reversion events observed in breast, ovarian, prostate and pancreatic cancer patients. Clinical, genomic, and histological information was retrieved and verified, and annotations standardized using HGVS nomenclature via Ensembl (release 114) and ANNOVAR. Key reversion features-including mutation consequence, location, proximity to the original mutation, domain involvement, and microhomology-were analyzed using in-house R scripts. **(B)** Reversion mutations in the *BRCA1/2* reversion cohort are predominantly reported in small studies. Bar chart showing the number of patients with reversion mutations reported per study and coloured by tumor type.

**Fig. S2.**
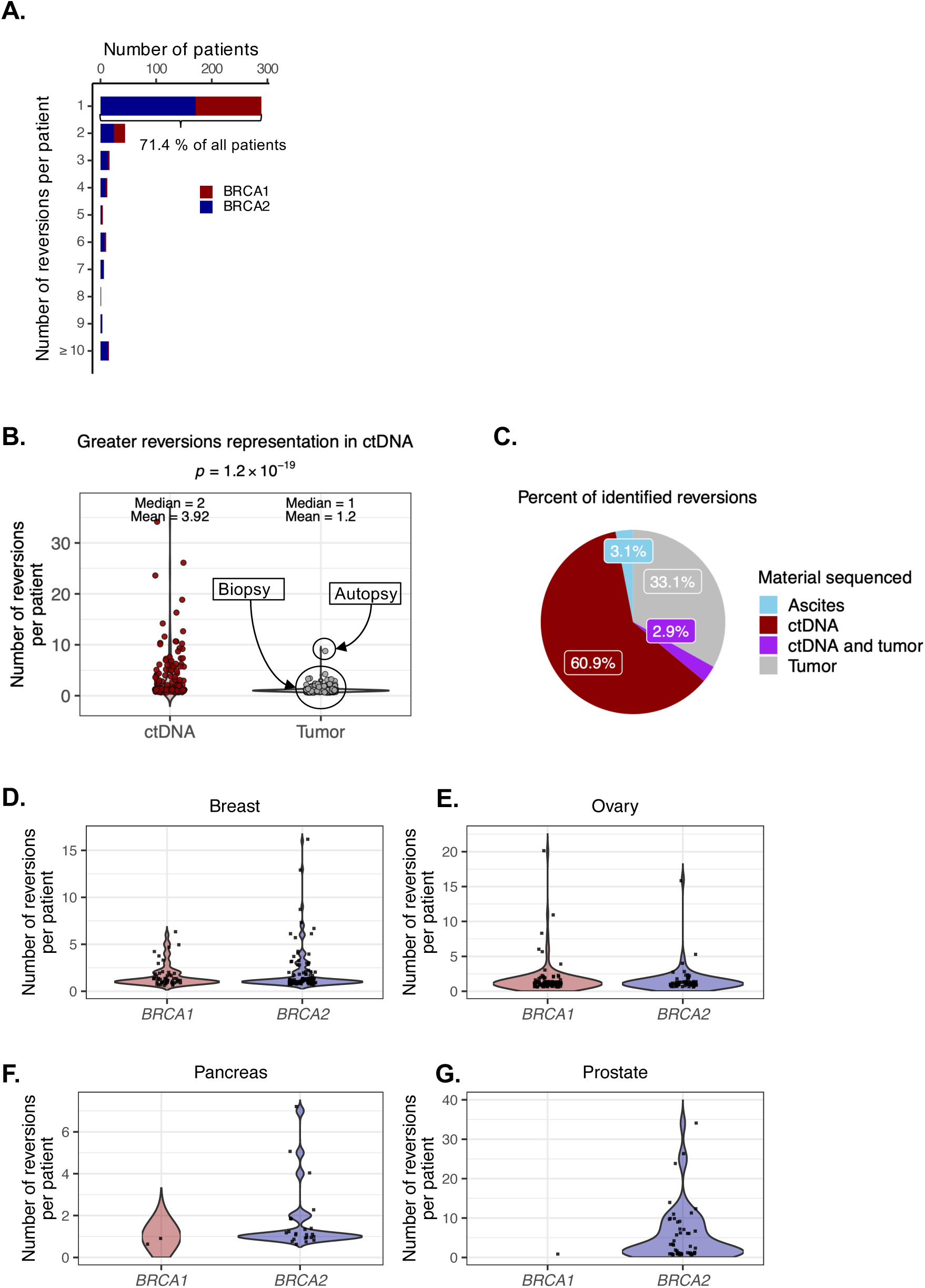
Reversion mutations per patient. **(A)** Column chart illustrating the number of reversion events per patient in *BRCA1* (red) and *BRCA2* (blue) mutant patients in the reversion cohort. Most patients (71.4 %) harbored a single reversion event. **(B)** Violin plots comparing the number of reversion mutations identified per patient detected in blood (ctDNA) versus solid tumor sequencing biopsies for *BRCA1* and *BRCA2* genes. Multiple reversion mutations in a single patient are more frequently observed in blood samples compared to tumor samples (Wilcoxon test *p* = 1.2 x 10^-19^). **(C)** Pie chart showing that the majority of the reversions annotated in the reversions cohort (60.9%) were detected in ctDNA. **(D-G)** Violin plots showing the number of different *BRCA1/2* reversion mutations detected per patient in breast, ovarian, pancreatic and prostate cancers. Prostate cancer displays some extreme outlier cases, including patients with up to 34 reversions in (*18*) and up to 26 reversions in (*23*) identified using reversion detection pipeline AARDVARK (*13*).

**Fig. S3.**
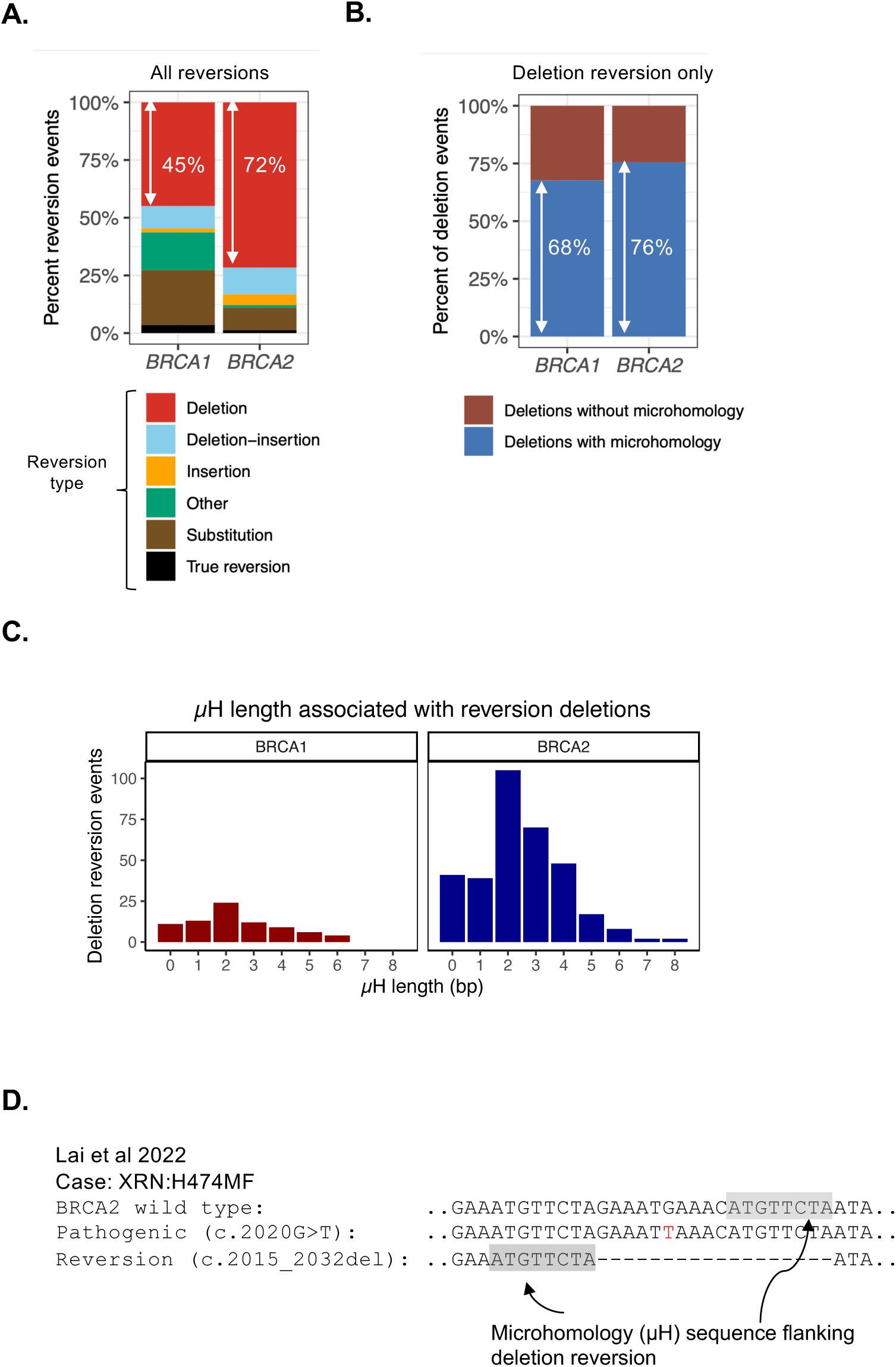
Microhomology in *BRCA1/2* reversion deletions. **(A)** Column chart summarizing reversion mutation types in *BRCA1* and *BRCA2* mutant patients, classified into deletions, deletion–insertions, insertions, substitutions, true reversions and other categories. Deletion reversions (either microhomology flanked or not) account for 45% *BRCA1* reversions and 72% of *BRCA2* reversions. **(B)** Column chart summarizing the relative frequency of microhomology-associated deletion reversions (68% of all deletion reversions in *BRCA1* and 76% of deletion reversions in *BRCA2*) compared to deletion reversions not associated with flanking microhomology. **(C)** Distribution of microhomology (µH) lengths observed in deletion reversion events across *BRCA1* (left) and *BRCA2* (right). The majority of reversion deletions were associated with short µH sequences (1-3 bp), with 2bp µH being the most frequently used length in both *BRCA1* and *BRCA2*. **(D)** Example of the longest µH length (8 bp in length) identified.

**Fig. S4.**
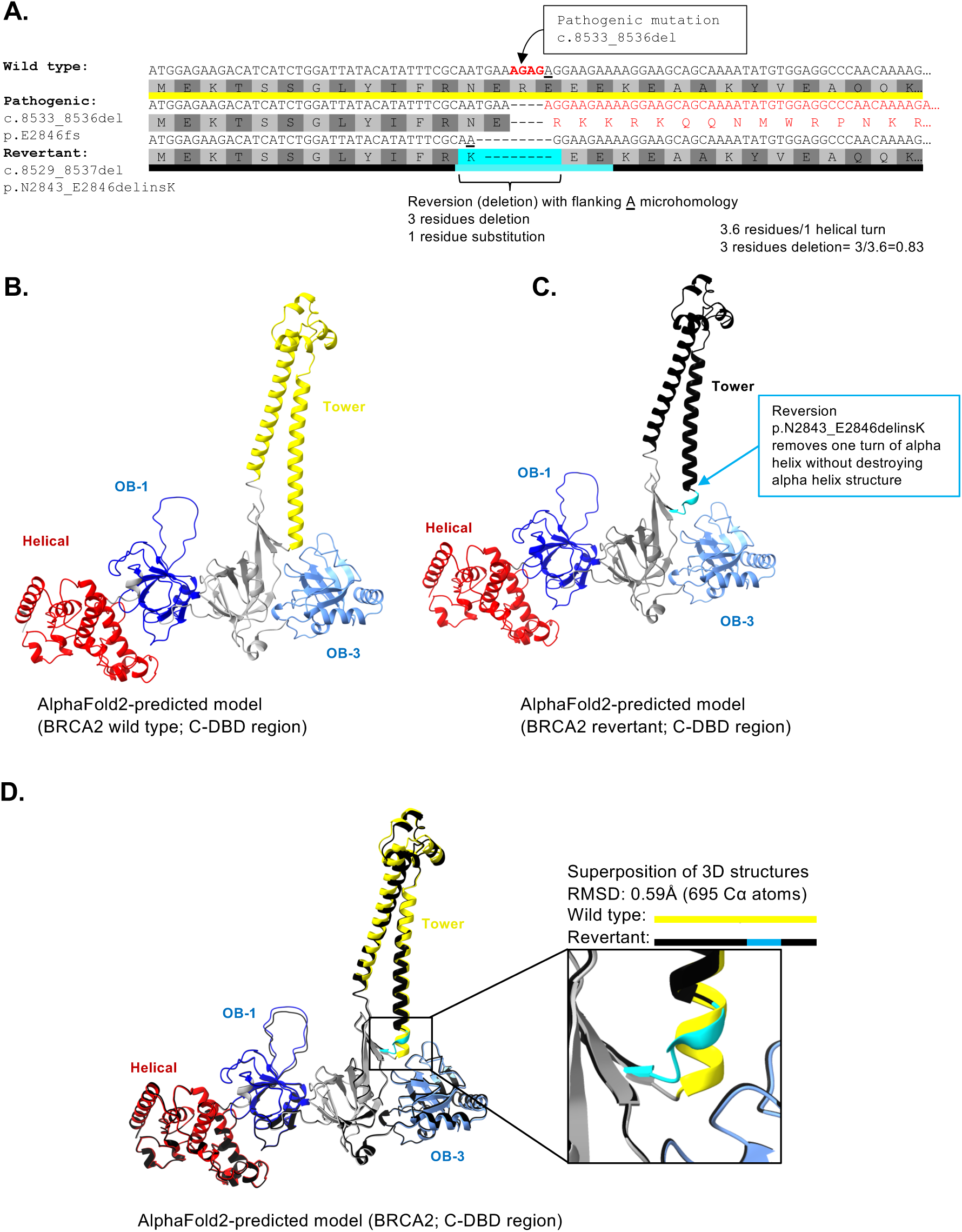
Minimal structural impact of a reversion in the BRCA2 tower domain, observed in a patient in ARIEL2 trial. **(A)** DNA and amino acid sequence alignment of the wild-type, pathogenic, and reverted *BRCA2* alleles, showing the local region surrounding the pathogenic mutation and the reversion event identified in subject 85 from the ARIEL2 trial. **(B)** Predicted structural model of the wild type BRCA2 C-DBD region highlighting the tower domain (yellow), adjacent OB-fold domains (OB-1 and OB-3) and helical domain (red). **(C)** Predicted structural model of the reverted BRCA2 C-DBD from (A) showing a truncated tower domain (black). The three amino acid deletion and one amino acid substitution in the reverted protein is predicted to remove one turn of the tower alpha helix without disrupting the global fold of the tower domain and adjacent α-helical region. **(D)** Superposition of wild-type and revertant models using UCSF Chimera X (*52*) shows minimal structural differences, with a low RMSD value (0.59 Å over 695 Cα atoms).

**Fig. S5.**
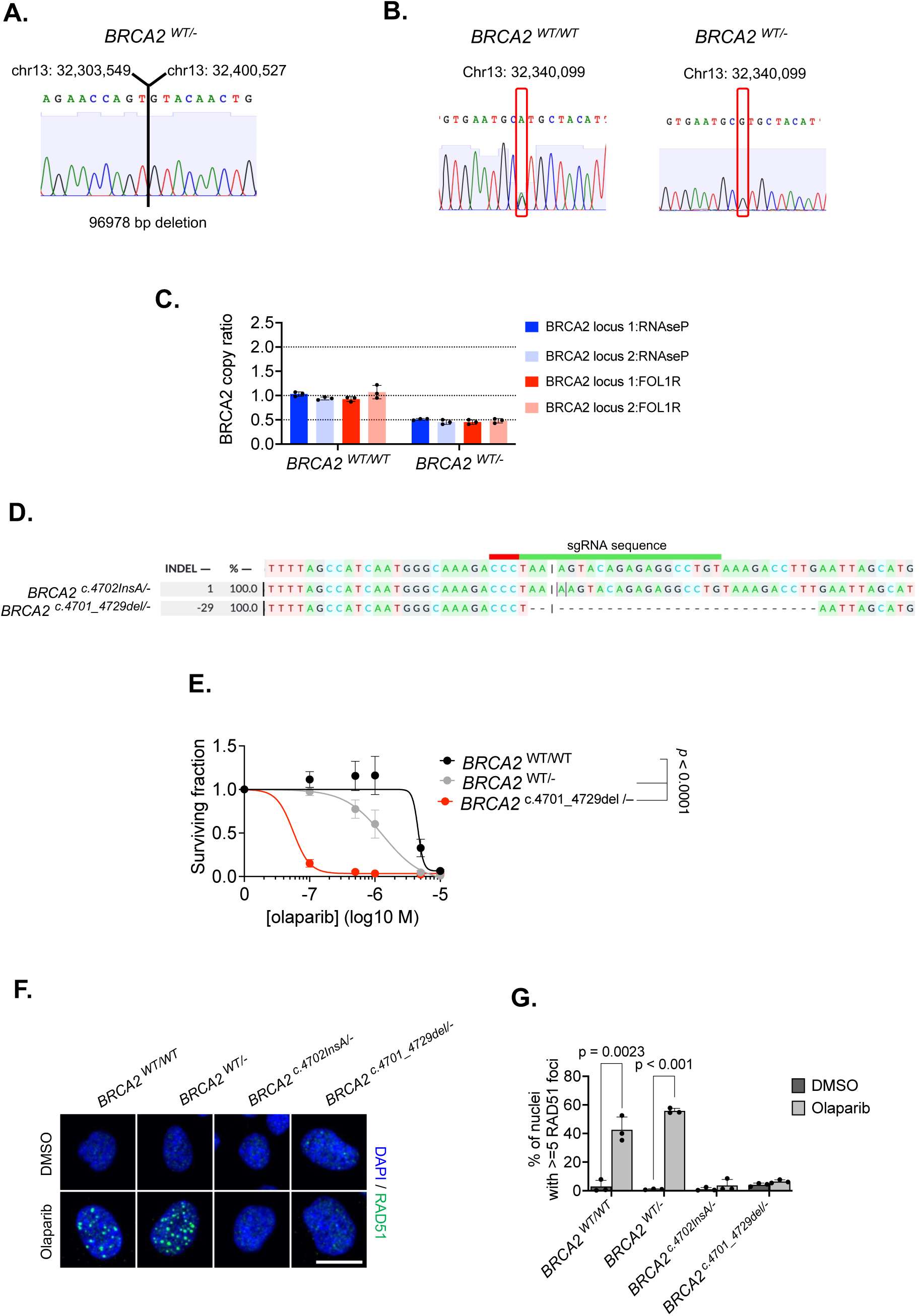
Generation of *BRCA2* mutant cells. **(A)** Sanger sequencing trace of the RPE1 *P53*^−/–^*BRCA2* ^WT/WT^ derived clone, *BRCA2* ^WT/–^. CRISPR-Cas9 mutagenesis was used to delete 96,978 bp of human chromosome 13 that encapsulated one copy of *BRCA2* (see Figure 6A); the sequencing trace across the resultant deletion breakpoint is shown with a vertical line defining the deletion breakpoint. Flanking genomic co-ordinates are shown. **(B)** Loss of one copy of BRCA2 in *BRCA2* ^WT/–^ cells was confirmed by genotyping of a heterozygous *BRCA2* single-nucleotide polymorphism (SNP). Sanger sequencing traces from *BRCA2* ^WT/WT^ and *BRCA2* ^WT/–^ cells are shown alongside the genomic coordinate of the SNP, which is heterozygous in *BRCA2* ^WT/WT^ cells and hemizygous in *BRCA2* ^WT/–^ cells. **(C)** Column chart illustrating digital droplet PCR (ddPCR) quantification of *BRCA2* copy number using *BRCA2*-specific ddPCR probes and diploid gene control loci (for *RNAseP* and *FOLR1*) in *BRCA2* ^WT/WT^ cells and *BRCA2* ^WT/–^ cells. Two different *BRCA2* loci were detected by ddPCR (locus 1 and locus 2) and the the ratio of positive droplets from *BRCA2* loci *vs.* control loci used to estimate *BRCA2* copy number in each sample. Data points show replica measurements. Error bars indicate standard deviation. **(D)** Sanger sequence DecodR (*53*) analysis of *P53^-/-^ BRCA2^WT/-^* derived clones *BRCA2* ^c.4702InsA/–^ and *BRCA2* ^c.4701_4729del/–^, after receiving the indicated crRNA sequence in exon11 of *BRCA2,* showing the number of base pairs gained/deleted (INDEL) relative to the unedited sequence and percentage each edit contributes to the sanger sequence trace (%). **(E)** PARPi dose-response curves in *BRCA2*^WT/WT^, *BRCA2* ^WT/–^ and *BRCA2* ^c.4701_4729del/–^ cells exposed to the PARPi olaparib for 10 days, after which survival was assessed using CellTiter Glo. Surviving fractions compared to cells exposed to the drug vehicle (DMSO) are shown. *p* values calculated using ANOVA. Error bars represent standard deviation from n = 3 experiments. *BRCA2*^WT/WT^ and *BRCA2* ^WT/–^ are also shown in Fig. 6B. **(F)** Representative images of nuclear RAD51 foci in *BRCA2*^WT/WT^, *BRCA2* ^WT/–^, *BRCA2* ^c.4702InsA/–^ and *BRCA2* ^c.4701_4729del/-^ cells exposed to the PARPi olaparib (5 μM) or the drug vehicle, DMSO, for 24 hours. Scale bar = 10 μm. Blue staining represents nuclei (DAPI), green staining represents RAD51. **(G)** Percentage of cells with >=5 nuclear RAD51 foci in cells exposed to either 5 µM olaparib (or drug vehicle, DMSO) for 24 hours. Error bars represent standard deviation from n = 3 experiments. *p* values calculated by Student’s t test.

**Fig. S6.**
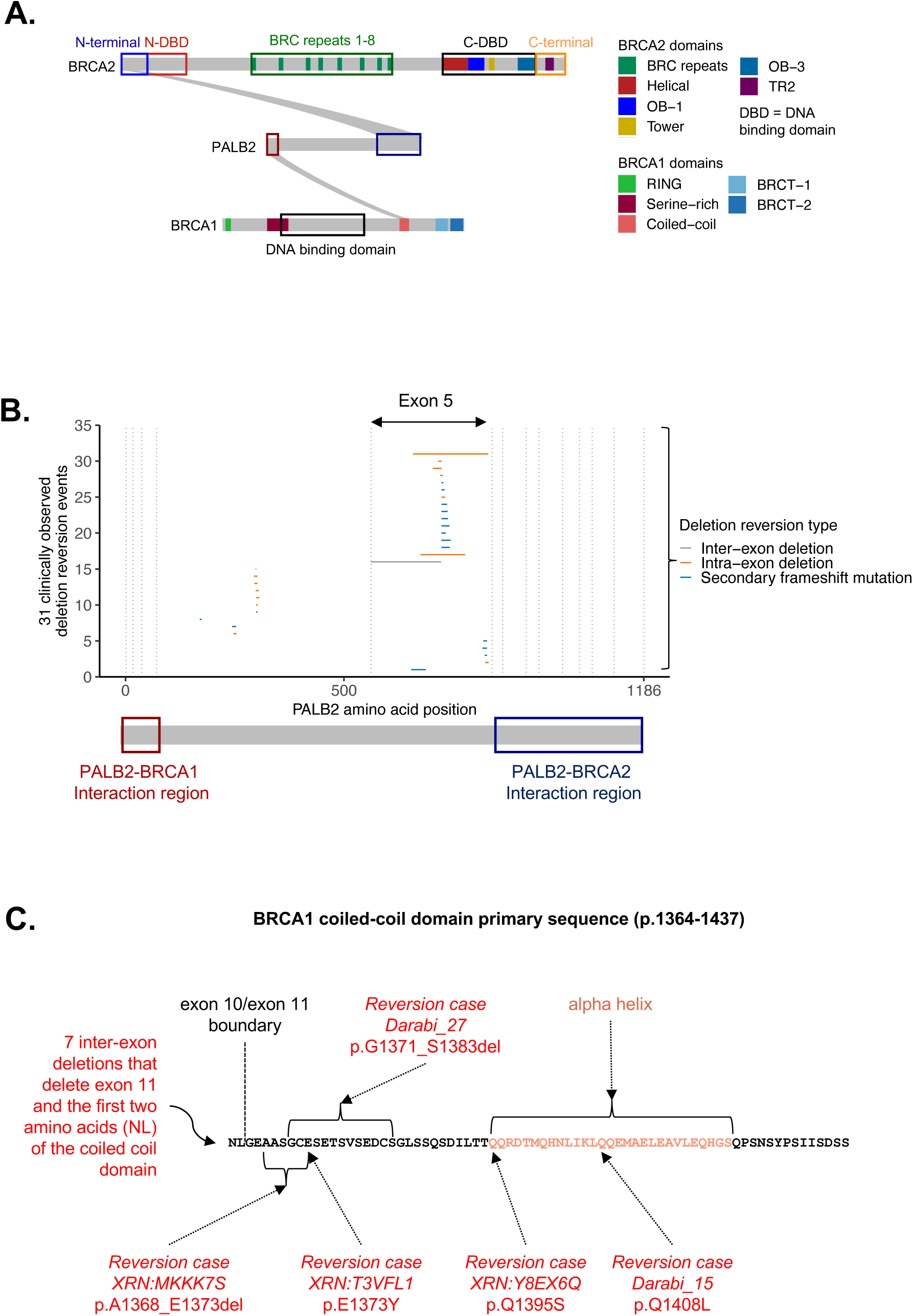
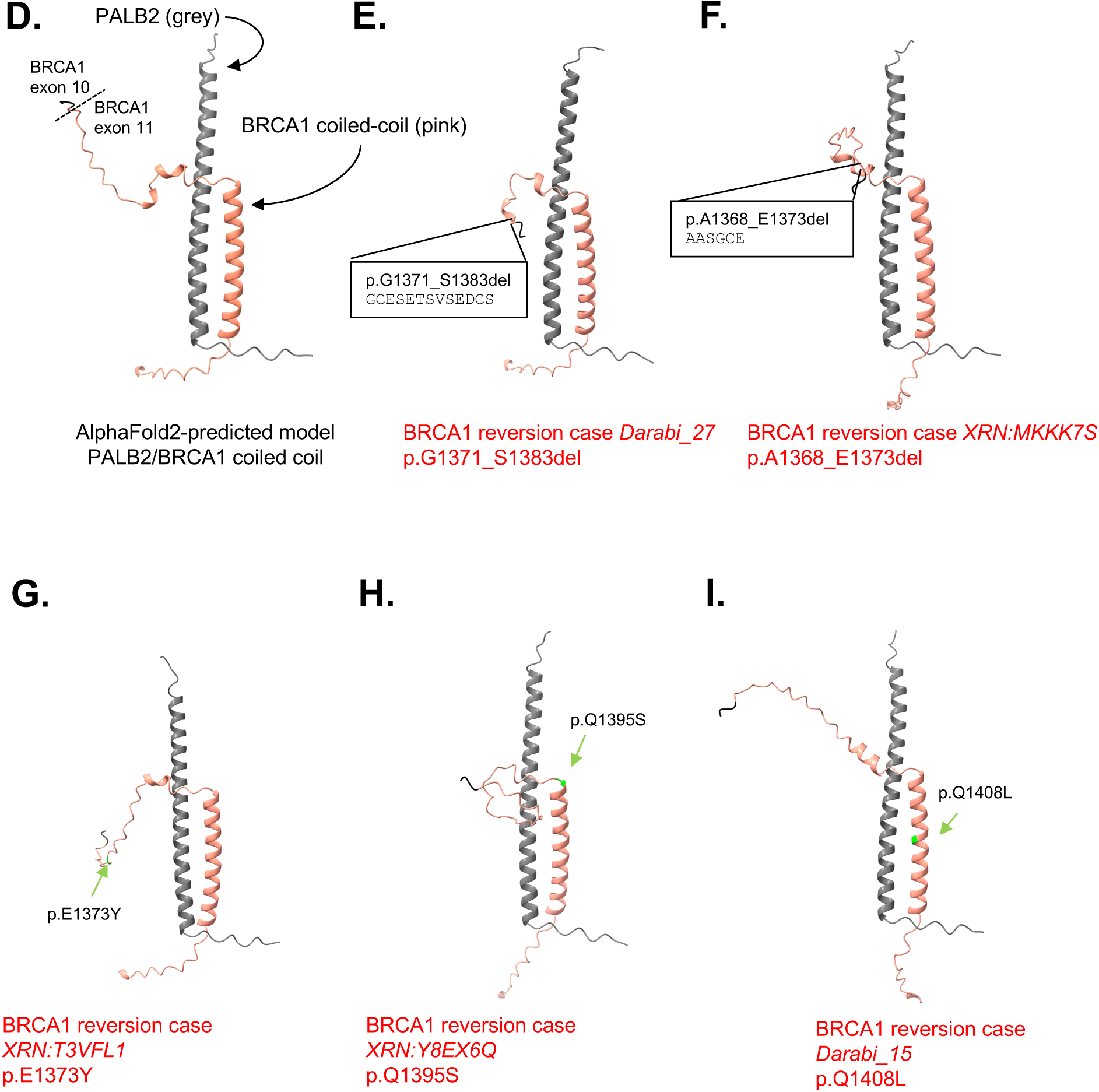
Reversions in PALB2 preserve BRCA1/2-interacting domains. **(A)** Schematic representation of PALB2 interactions with BRCA1 and BRCA2 proteins. Domain structures of BRCA1 and BRCA2 proteins are shown, highlighting the BRCA2 N terminal and BRCA1 coiled coiled coil domains that facilitate interactions with the PALB2 C-terminus (for BRCA2) and PALB2 N-terminus (for the BRCA1 interaction). **(B)** Alignment of *PALB2* deletion reversions with PALB2 amino acid positions. Deletions (horizontal lines) depict the three types of deletion reversions: Inter-exon deletion (grey), intra-exon deletion (orange), and secondary frameshift mutations (blue). Vertical lines indicate exon boundaries. Below the plot, PALB2 protein domains are displayed. **(C)** Primary amino acid sequence of the BRCA1 coiled coil domain that interacts with PALB2. Within the domain is an alpha helix (depicted in pink) that interacts with a complementary alpha helix in PALB2. Shown are the positions of 12 BRCA1 reversions that alter the primary sequence of the BRCA1 coiled coil domain; seven were deletions in exon 10 affecting only the first two amino acids and thus predicted to encode the complete BRCA1 alpha helix. The other reversions (annotated in red, with case numbers in *italics*) either altered a disordered region of BRCA1 N-terminal to the alpha helix, without altering the structure of the alpha helix itself (p.A1368_E1373del, p.G1371_S1383del or p.E1373Y) or changed the primary sequence of the alpha helix without obviously disturbing its tertiary structure (p.Q1395S, p.Q1408L). **(D-I)** AlphaFold2-predicted (*54*) structural models of the PALB2/BRCA1-coiled-coil complex, including the wild-type interaction **(D)** and models for each of the BRCA1 reversions described in **(C)**. For **(I)**, the model of PALB2 and BRCA1 p.Q1408L, the stability calculation for BRCA1 p.Q1408L (using FoldX (*55*)) indicated a negligible energetic impact (ΔΔG = −0.06 kcal/mol), suggesting that this substitution is unlikely to significantly alter the stability of the BRCA1-PALB2 complex.

**Fig. S7.**
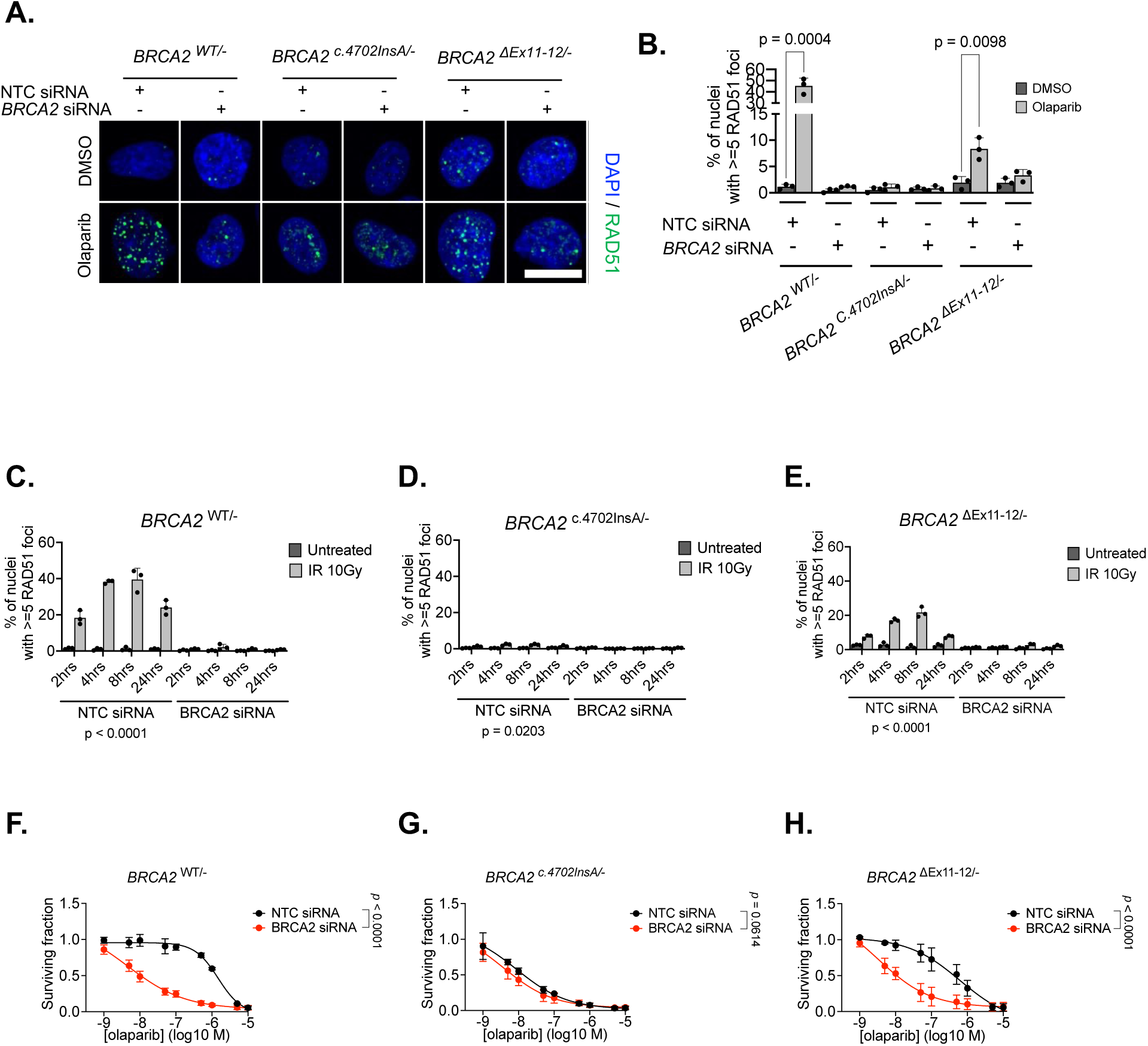
Cells lacking exons 11-12 of BRCA2 show restoration of RAD51 foci and PARPi resistance. **(A)** Representative images of nuclear RAD51 foci in *BRCA2*^WT/-^, *BRCA2*^c.4702InsA/–^ and *BRCA2*^ΔEx11-12/-^ cells exposed to the PARPi olaparib (5 μM) or the drug vehicle, DMSO, for 24 hours following exposure to non-targeting control (NTC) or BRCA2 siRNA. Scale bar = 10 μm. Blue staining represents nuclei (DAPI), green staining represents RAD51. **(B)** Percentage of cells with >=5 nuclear RAD51 foci in cells exposed to either 5 µM olaparib (or drug vehicle, DMSO) for 24 hours following exposure to NTC or BRCA2 siRNA. Error bars represent standard deviation from n = 3 experiments. *p* values calculated by Student’s t test. **(C-E)** Percentage of cells with >=5 nuclear RAD51 foci in *BRCA2*^WT/-^ (C), *BRCA2* ^c.4702InsA/–^ (D) and *BRCA2* ^ΔEx11-12/–^ (E) cells at the indicated time point after receiving ionizing radiation (10 Gy) or without treatment, following exposure to NTC or BRCA2 siRNA. Error bars represent standard deviation from n = 3 experiments. *p* values calculated via two-way ANOVA. **(F-H)** PARPi dose-response curves in *BRCA2*^WT/-^ (F), *BRCA2* ^c.4702InsA/–^ (G) and *BRCA2* ^ΔEx11-12/–^ (H) cells exposed to the PARPi olaparib for 6 days following exposure to NTC or BRCA2 siRNA, after which survival was assessed using CellTiter Glo. Surviving fractions compared to cells exposed to the drug vehicle (DMSO) are shown. *p* values calculated using ANOVA. Error bars represent standard deviation from n = 3 experiments.

**Fig. S8.**
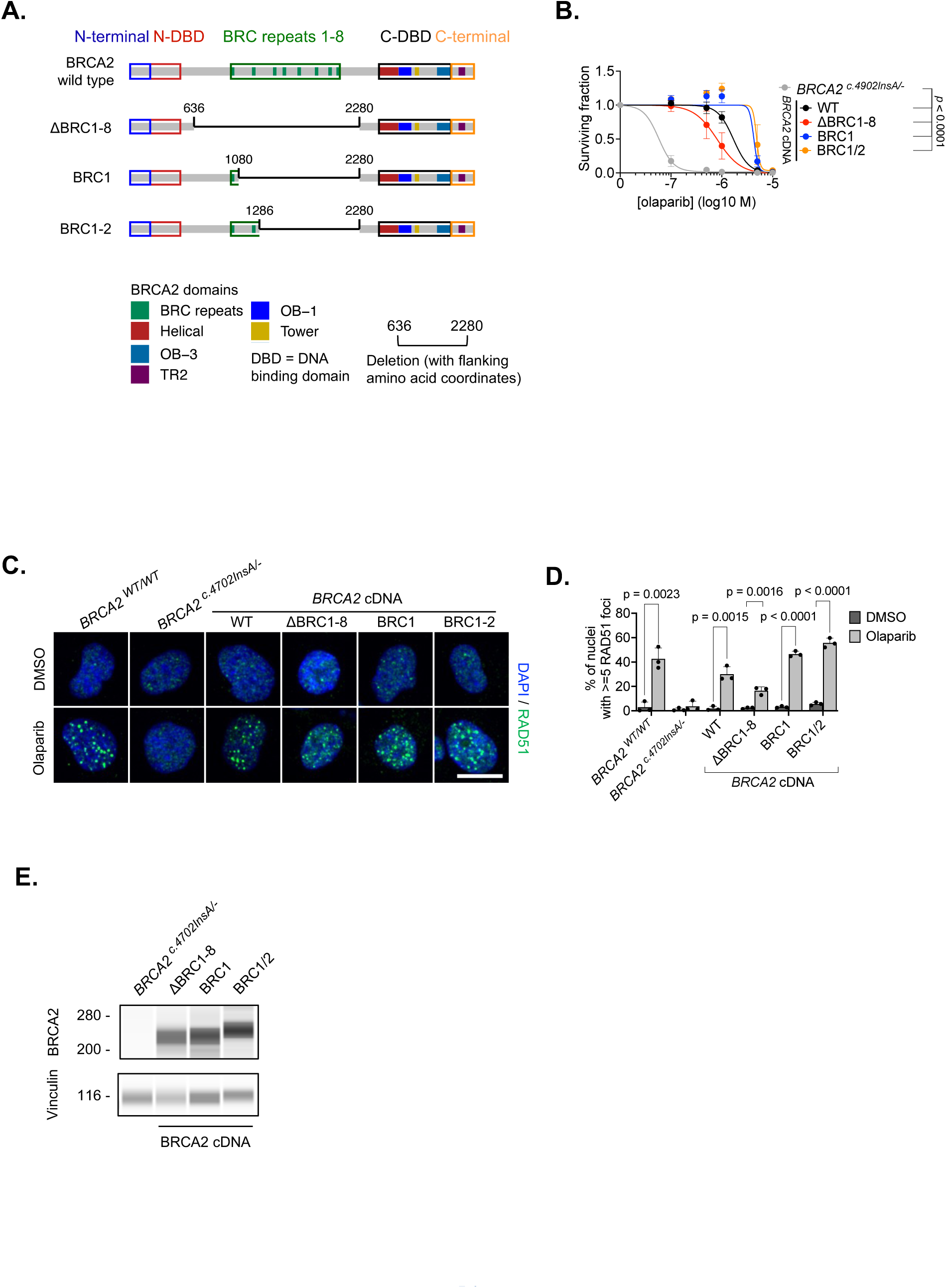
*BRCA2* transgene variants lacking BRC repeats confers PARPi resistance and RAD51 foci. **(A)** Graphical representation of BRCA2 protein variants expressed via cDNA constructs. ΔBRC1-8, BRC1 and BRC1/2, encode progressive additions of BRC domains, starting from deletion of all BRC repeats (ΔBRC1-8). Deleted regions are indicated with a thin line and flanking amino acid co-ordinates. **(B)** PARPi dose-response curves in *BRCA2* ^c.4702InsA/–^ cells expressing cDNAs from (A) exposed to the PARPi olaparib for 10 days, after which survival was assessed using CellTiter Glo. Surviving fractions compared to cells exposed to the drug vehicle (DMSO) are shown. *p* values calculated using ANOVA. Error bars represent standard deviation from n = 3 experiments. **(C)** Representative images of nuclear RAD51 foci in *BRCA2* ^c.4702InsA/–^cells expressing cDNAs from (A) exposed to the PARPi olaparib (5 μM) or the drug vehicle, DMSO, for 24 hours. Scale bar = 10 μm. Blue staining represents nuclei (DAPI), green staining represents RAD51. **(D)** Percentage of cells with >=5 nuclear RAD51 foci in cells exposed to either 5 µM olaparib (or drug vehicle, DMSO) for 24 hours. Error bars represent standard deviation from n = 3 experiments. *p* values calculated by Student’s t test. (**E**) Simple Western immunoblot analysis using an antibody detecting a BRCA2 C-terminal epitope in lysates generated from BRCA2 c.4702InsA/– cells expressing BRCA2 cDNA constructs. Positions of molecular weight markers shown on left (kDa).

**Fig. S9.**
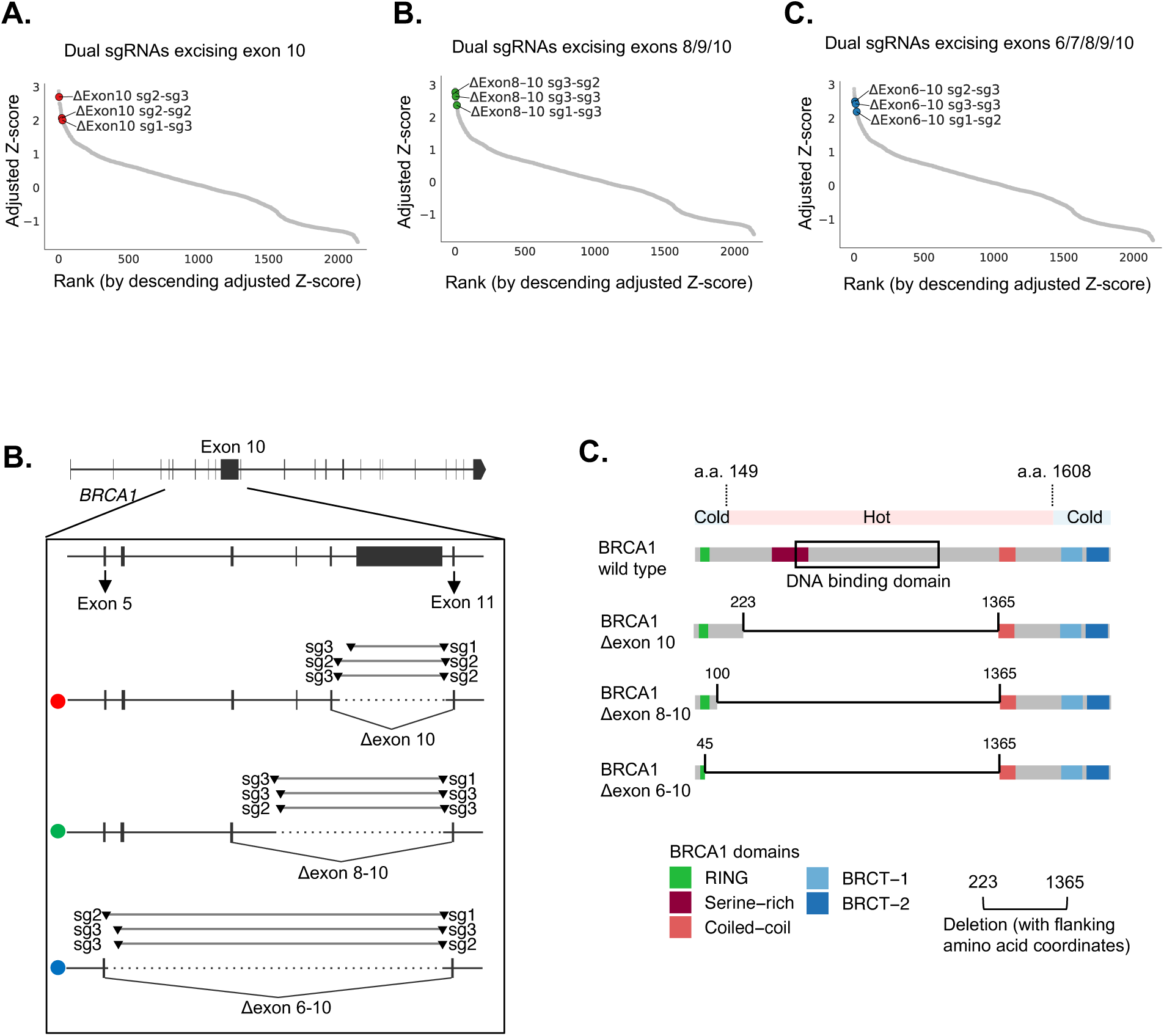
BRCA1 exon excision CRISPR screening identifies BRCA1 regions dispensable for PARPi resistance. (**A**) Adjusted Z-score of *BRCA1* CRISPR sgRNA pair read counts, ranked in descending order following olaparib exposure. Vectors for intron combinations with at least three vectors with adjusted Z > 2 are highlighted. (**B**) Representation of *BRCA1* intron targets for highlighted sgRNA pairs which had an adjusted Z-score of >2, showing theoretical exon excision events. (**C**) Schematic of predicted BRCA1 proteins resulting from the excision of indicated exons, compared to wild-type (WT) and *BRCA1* reversion hot and cold regions.

**Fig. S10.**
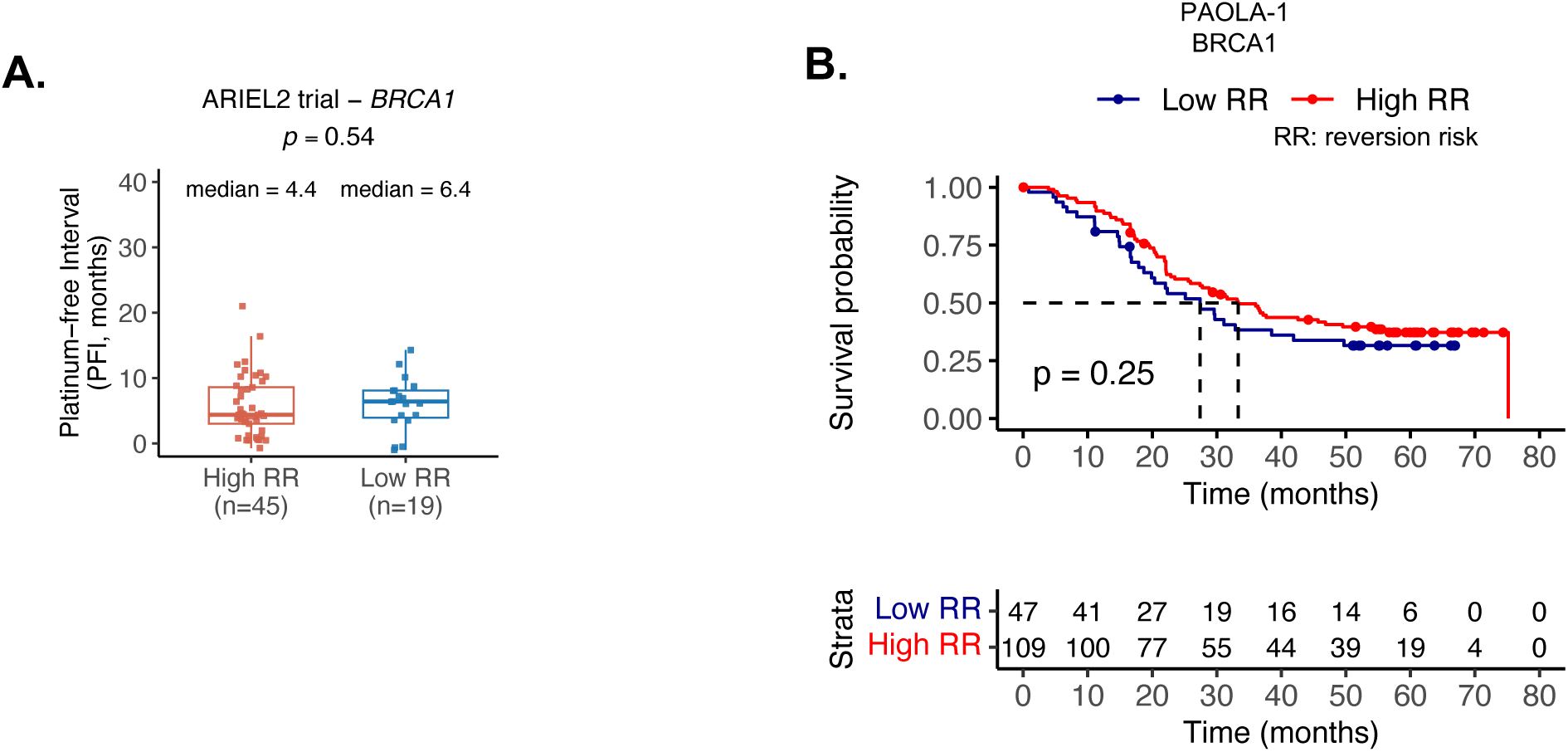
**(A)** Lack of correlation between *BRCA1* reversion risk and PFI in ARIEL2 trial. Patients were stratified into high (red) or low (blue) reversion-risk groups. *p* value calculated with Wilcoxon test. **(B)** Lack of correlation between *BRCA1* reversion risk and PFS in PAOLA-1 trial. Patients were stratified into high (red) or low (blue) reversion-risk groups. *p* values were calculated using the log-rank test.

Data S1. Literature used to collate the reversions cohort

Data S2. Pathogenic BRCA1/2 mutations in the reversion cohort

Data S3. Pathogenic BRCA1/2 mutations and identified reversion mutations in the reversion cohort

Data S4: BRCA1/2 reversion mutation types in the reversions cohort

Data S5: BRCA1/2 reversion deletion types in BRCA1/2 in the reversion cohort

Data S6. BRCA1/2 pathogenic mutation types in the reversions cohort

Data S7: Incidence cohort.

Data S8: BRCA1/2 pathogenic mutation types in the incidence cohort

Data S9: Comparison of mutation type frequencies between reversion and incidence cohorts

Data S10. Pathogenic PALB2 mutations with identified clinical reversions

Data S11. Pathogenic PALB2 mutations and the identified reversion mutations

Data S12: The Incidence cohort mutation classification list

Data S13: The BTBC cohort mutation classification list

Data S14: ARIEL2 trial mutation classification list

Data S15: F1 scores associated with reversion predictor

Data S16: PAOLA-1 trial mutation classification list

Data S17. Nucleic acid sequences used

Data S18: sgRNA sequences used in dual CRISPR screens

